# Endogenous RALF peptide function is required for powdery mildew host colonization

**DOI:** 10.1101/2025.01.30.635691

**Authors:** Henriette Leicher, Sebastian Schade, Jan W. Huebbers, Kristina S. Munzert-Eberlein, Genc Haljiti, David Biermann, Athanasios Makris, Xiaoxuan Zhu, Yashank Chauhan, Christina Ludwig, Marion C. Müller, Toshinori Kinoshita, Timo Engelsdorf, Julien Gronnier, Martina K. Ried-Lasi, Aurélien Boisson-Dernier, Ralph Hückelhoven, Martin Stegmann

## Abstract

The receptor kinase FERONIA (FER) is a susceptibility factor for biotrophic powdery mildew fungal pathogens in *Arabidopsis thaliana*, but the underlying molecular mechanisms remain largely unknown. FER is required for the perception of endogenous RAPID ALKALINIZATION FACTOR (RALF) peptides to control various aspects of plant growth, development and immunity. RALFs are perceived by FER-LORELEI-LIKE GPI-ANCHORED PROTEIN (LLG) heterocomplexes to induce cellular responses and bind to LEUCINE-RICH REPEAT EXTENSIN (LRX) proteins as structural components of the cell wall. Combining genetics, cell biology and biochemistry, we found that FER’s endogenous RALF ligands are necessary for full colonization success of the powdery mildew species *Erysiphe cruciferarum*. We reveal that LLGs and LRXs are also powdery mildew susceptibility factors. We show that cell wall remodeling and apoplastic pH homeostasis, hallmark features of RALF function, support powdery mildew reproductive success. Moreover, we provide data that RALF-dependent powdery mildew pathogenesis is partially independent of FER. Powdery mildew fungi likely do not produce RALF peptide mimics, suggesting their reliance on endogenous RALFs for successful host colonization. We propose that powdery mildew fungi require RALF-mediated modulation of apoplastic pH and pectin re-modelling for successful host colonization, highlighting a new susceptibility mechanism by obligate biotrophic fungi.

## Introduction

Powdery mildew is one of the most widespread fungal plant diseases affecting several economically relevant crop species (Glawe, 2008), with infection being characterized by continuous interaction between the biotrophic fungus and its host plant. Susceptibility genes (S-genes) can affect fungal infection success and are utilized by pathogens for their own benefit. In consequence, mutating these genes usually confers pathogen and genotype-independent and hence durable disease resistance (van Schie & Takken, 2014; Engelhardt *et al*., 2018). The first well-characterized powdery mildew susceptibility factor was the barley MILDEW LOCUS O (MLO) that confers barley susceptibility towards adapted *Blumeria hordei*. Mutants in orthologous *MLO* genes are powdery mildew resistant in phylogenetically diverse plant species, including *Arabidopsis thaliana* (hereafter Arabidopsis) and tomato (Kusch & Panstruga, 2017).

Next to susceptibility factors, the outcome of plant-pathogen interactions is shaped by their immune system. Pattern-triggered immunity (PTI) is the first line of induced resistance and activated by the perception of microbe-associated molecular patterns (MAMPs) by pattern recognition receptors (PRRs), which are often plasma membrane localized receptor-like kinases (RLKs). To subvert PTI, adapted pathogens evolved effectors that are mainly perceived by intracellular nucleotide-binding leucine-rich repeat receptors (NB-LRRs) activating pathogen race-specific effector triggered immunity (Ngou *et al*., 2022).

In addition to their function as PRRs, RLKs are important sensors for endogenous signals regulating growth, abiotic stress and/or modulating immunity (Rzemieniewski & Stegmann, 2022; Zecua-Ramirez *et al*., 2026). Adapted pathogens can hijack RLK pathways to support accommodation (Hok *et al*., 2011; Hok *et al*., 2014; Ried *et al*., 2019). The conserved RLK FERONIA (FER) and related CATHARANTUS ROSEUS RECEPTOR-LIKE KINASE 1-LIKEs (CrRLK1Ls) perceive endogenous RAPID ALKALINIZATION FACTOR (RALF) peptides to regulate multiple plant physiological processes (Ge *et al*., 2017; Gonneau *et al*., 2018; Zhu *et al*., 2021; Zhong *et al*., 2022; Lan *et al*., 2023). FER recognizes RALFs in a heteromeric complex with LORELEI-LIKE GPI ANCHORED PROTEIN 1 (LLG1) (Li *et al*., 2015; Xiao *et al*., 2019) and controls PTI as a RALF-regulated scaffold for PRR complexes and their nanoscale organization (Stegmann *et al*., 2017; Gronnier *et al*., 2022). In addition to FER-dependent signaling, recent reports show that RALFs have a structural role in organizing the patterning of demethylesterified pectins in the cell wall (Moussu *et al*., 2023; Schoenaers *et al*., 2024). Here, RALFs bind to cell wall bound LEUCINE-RICH REPEAT EXTENSIN (LRX) proteins but the biochemical interplay between LRX and FER-LLG complexes remains little understood (Mecchia *et al*., 2017; Zhao *et al*., 2018; Dünser *et al*., 2019; Moussu *et al*., 2020; Gronnier *et al*., 2022; Schade *et al*., 2025). Adapted pathogens can hijack RALF signaling for their own benefit. *Fusarium oxysporum* and root knot nematodes produce RALF mimics to enhance infection success on tomato and Arabidopsis (Masachis *et al*., 2016; Thynne *et al*., 2017; Zhang *et al*., 2020). Furthermore, the Ler-0 *fer-1* mutant is more resistant to the powdery mildew species *Golovinomyces orontii*, suggesting that FER acts as an *S*-factor (Kessler *et al*., 2010). Yet, how FER confers powdery mildew susceptibility remains largely unknown.

Here, we investigated the molecular mechanism of FER-mediated powdery mildew susceptibility, using the Arabidopsis-adapted powdery mildew fungus *Erysiphe cruciferarum* (*Ecr)*. We show that LLG1 and LRX1-LRX5 are required for successful powdery mildew infection. By loss of function studies, we identify RALF peptides as important susceptibility factors for powdery mildew. CRISPR Cas9-generated *ralf22/23/33* (CRISPR *ralf3x*) and *ralf1/18/22/23/31/33/34 (*CRISPR *ralf7x)* mutants show increasing resistance to *Ecr* infection depending on the number of eliminated RALFs. Our bioinformatic analysis indicates that powdery mildew fungi do not encode functional RALF peptide mimics, supporting that they require host-endogenous RALFs for their own benefit (Masachis *et al*., 2016; Thynne *et al*., 2017). We suggest that the enhanced resistance of *fer* and CRISPR *ralf7x* is not associated with defects in MLO-mediated penetration, or deregulation of defense-related phytohormone signaling. Instead, powdery mildew might require FER-RALF-mediated apoplastic pH regulation and cell wall remodeling for successful colonization. Surprisingly, our data indicates that RALF-mediated powdery mildew susceptibility is partially independent on FER and exerts its effect in a combination of signaling and structural functions. We propose that powdery mildew fungi require RALFs to provide a favorable growth environment, highlighting a new mechanism of host subversion by adapted plant pathogens.

## Results

### FER, LLGs and LRX are powdery mildew susceptibility factors

Ler-0 *fer-1* is more resistant to *Golovinomyces orontii* (Kessler *et al*., 2010). We confirmed this phenotype in Arabidopsis Col-0 using the *fer-4* mutant allele (Duan *et al*., 2010) and *Ecr* (Micali *et al*., 2008). The *fer-4* mutant showed slightly weaker visible symptoms (Fig. 1A), reduced conidiophore production and less relative fungal DNA content compared to Col-0 5 dpi (Fig. 1B, C, Fig. S1A). Expression of pFER::FER-GFP complemented the *fer-4* resistance phenotype (Fig. 1C).

**Figure 1:**
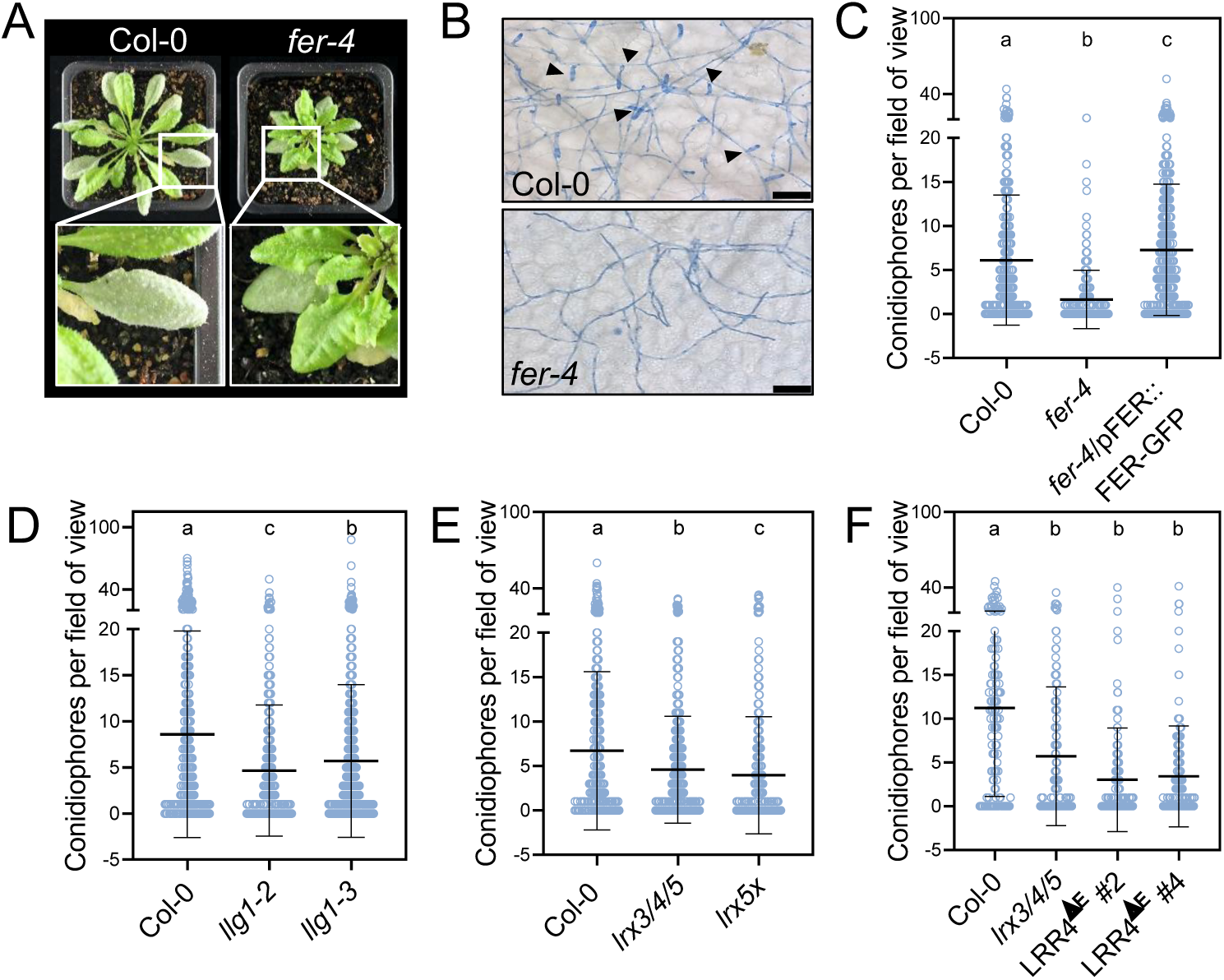
**FER and additional RALF binding proteins are powdery mildew susceptibility factors.** A) Images of Col-0 and *fer-4* plants infected with *Ecr* (14 dpi). B) Ink stained *Ecr* colonies (5 dpi) grown on Col-0 and *fer-4* plants. Black arrows indicate conidiophores. Scale bar represents 50 µm. C) Conidiophores per field of view (5 dpi) of fungal colonies grown upon *Ecr* infection of the indicated genotypes. Mean ± SD, n=223-392 pooled from three independent experiments (Dunn’s multiple comparisons test, a-b, b-c p<0.0001; a-c p=0.0106). D) Conidiophores per field of view (5 dpi) of fungal colonies grown upon *Ecr* infection of the indicated genotypes. Mean ± SD, n=345-665 pooled from five independent experiments (Dunn’s multiple comparisons test, a-b, a-c p<0.0001; b-c p=0.0235). E) Conidiophores per field of view (5 dpi) of fungal colonies grown upon *Ecr* infection of the indicated genotypes. Mean ± SD, n=285-453 pooled from three independent experiments (Dunn’s multiple comparisons test, a-b p=0.0150; a-c p<0.0001; b-c p=0.0408). F) Conidiophores per field of view (5 dpi) of fungal colonies grown upon *Ecr* infection of the indicated genotypes (LRR4^ΔE^: LRX4 lacking the extensin domain). Mean ± SD, n=113-159 pooled from two independent experiments (Dunn’s multiple comparisons test, a-b p<0.0001). All experiments were done at least three times with similar results, except F), which was repeated once with identical results.

Next, we investigated whether FER-related RALF receptor complex components are involved in *Ecr* susceptibility. The *llg1-2* mutant showed reduced conidiophore production (Fig. 1D). A similar effect was observed in the *llg1-3* allele, in which LLG1 is impaired in RALF binding but mutants do not show growth phenotypes (Fig. 1D, Fig. S11) (Shen *et al*., 2017; Xiao *et al*., 2019). The resistance phenotype of the *llg1* mutants, however, was not clear in qPCR quantification (Fig. S1B), indicating potential redundancy of LLG1 with related members of the LRE/LLG gene family (Xiao *et al*., 2019; Noble *et al*., 2022). Surprisingly, *llg1-2* and *llg1-3* were previously described as more susceptible to *Golonivomyces cichoracearum* (Shen *et al*., 2017), raising the question of mildew-species-dependent differences in *llg1* pathophenotypes, unlike *fer* (Fig. 1C, D) (Kessler *et al*., 2010). RALFs also bind to cell wall localized LRX proteins (Moussu *et al*., 2020). A *lrx3 lrx4 lrx5* (*lrx3/4/5*) triple mutant phenocopies *fer* in immune signaling, salt stress responses and plant morphology (Zhao *et al*., 2018; Dünser *et al*., 2019; Moussu *et al*., 2020; Gronnier *et al*., 2022). This *lrx3/4/5* mutant displayed reduced conidiophore production upon *Ecr* infection (Fig. 1E). Yet, relative fungal DNA content was unaltered (Fig. S1C). Further mutating *LRX1* and *LRX2* in *lrx3/4/5* (*lrx1/2/3/4/5*, hereafter *lrx5x*) (Herger *et al*., 2020) promoted the resistance phenotype (Fig. 1E, Fig. S1C). Overexpression of LRR4-citrine (LRR4^ΔE^/LRX4 lacking the extensin domain) has a dominant negative effect on plant growth, while PTI responses are not affected (Dünser *et al*., 2019; Gronnier *et al*., 2022). The p35S::LRR4-citrine lines phenocopied *lrx5x* and *fer*-*4* in showing reduced *Ecr* conidiophore production (Fig. 1F).

### Powdery mildew requires endogenous RALF peptides for successful colonization

We hypothesized that RALF signaling through FER-LLG1 and/or their binding to LRXs is required for *Ecr* colonization. Many plant pathogens secrete RALF peptide mimics to promote virulence (Masachis *et al*., 2016; Thynne *et al*., 2017; Zhang *et al*., 2020). We searched available genomes of powdery mildew species for the presence of potential RALFs. However, consistent with previous reports (Thynne *et al*., 2017) we could not detect evidence for RALFs among different *Erysiphales* species (Table S1, method section for details on our bioinformatic search pipeline). Thus, we hypothesized that endogenous RALFs are involved in the *Ecr-*Arabidopsis interaction. Arabidopsis encodes for 37 RALFs (Abarca *et al*., 2021). We anticipated genetic redundancy and generated CRISPR mutants in which we targeted *RALF23*, a described FER-LLG1 ligand (Stegmann *et al*., 2017; Xiao *et al*., 2019; Tang, J *et al*., 2022) and seven additional RALF-peptides with predicted leaf expression (*RALF1*, *RALF14*, *RALF18*, *RALF22*, *RALF23*, *RALF31*, *RALF33*, *RALF34*) (Fig. S2A). We could not obtain an octuple mutant but generated two higher order mutants, *ralf22/23/33* (CRISPR *ralf3x*) and *ralf1/18/22/23/31/33/34* (CRISPR *ralf7x*) (Fig. 2A, Fig. S2B, C). CRISPR *ralf3x* displayed a mild growth phenotype with slightly shortened petioles, while CRISPR *ralf7x* was strongly impaired in shoot growth (Fig. 2B, Fig. S2D, S11). Compared to *fer-4*, however, CRISPR *ralf7x* showed less pronounced shoot growth defects with a trend towards longer petioles (Fig. S2D) and an overall more wildtype-like morphology (Fig. 2B, Fig. S11). *Ecr* infection was reduced on both CRISPR *ralf3x* and CRISPR *ralf7x* (Fig. 2B-E). Notably, CRISPR *ralf7x* displayed slightly elevated resistance compared to *fer-4* (Fig. 2D). Expression of *p35S::RALF23* complemented CRISPR *ralf7x* growth and *Ecr* infection phenotypes (Fig. 2F, Fig. S2E, F, S11). Additionally, the previously reported *ralf1/22/23/33* quadruple mutant (Lan *et al*., 2023) was more resistant to *Ecr* with slightly less condiophore production compared to *fer-4* (Fig. 2A, G). We analyzed expression of the seven *RALFs* mutated in CRISPR *ralf7x* in Col-0 upon *Ecr* infection by qPCR. *RALF18* and *RALF23* were significantly upregulated at 4 and 5 dpi (Fig. S3). Together with our genetic data, this underlines the role of endogenous RALF peptides in *Ecr* host colonization.

**Figure 2:**
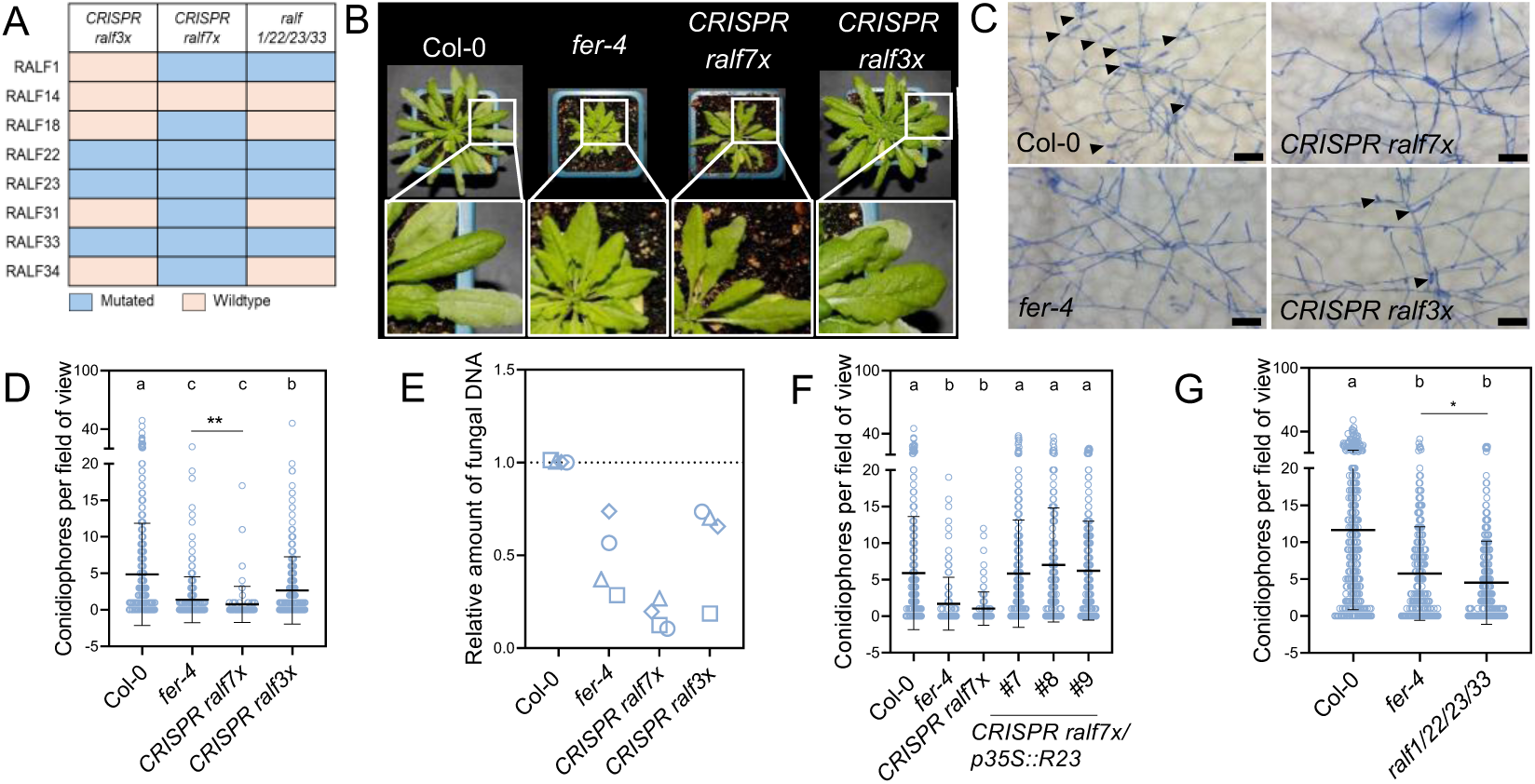
Specific RALFs are powdery mildew susceptibility factors. A) Table of RALF peptides mutated in CRISPR *ralf3x*, CRISPR *ralf7x* and *ralf1/22/23/33.* B) Images of Col-0*, fer-4*, CRISPR *ralf3x* and CRISPR *ralf7x* plants infected with *Ecr* (14 dpi). C) Ink stained *Ecr* colonies (5 dpi) grown on Col-0, *fer*-*4*, CRISPR *ralf3x* and CRISPR *ralf7x* plants. Black arrows indicate conidiophores. Scale bar represents 50 µm. D) Conidiophores per field of view (5 dpi) of fungal colonies grown upon *Ecr* infection of the indicated genotypes. Mean ± SD, n=98-456 pooled from three independent experiments (Dunn’s multiple comparisons test, a-b, a-c p<0.0001, b-c p<0.0002. Comparison between *fer-4* and CRISPR *ralf7x*: Mann Whitney test, ** p=0.007). E) Amount of fungal DNA normalized to plant DNA (5 dpi) upon *Ecr* infection of the indicated genotypes, n=4, data points with different symbols indicate independent biological replicates. F) Conidiophores per field of view (5 dpi) of fungal colonies grown upon *Ecr* infection of the indicated genotypes. Mean ± SD, n=101-286 pooled from three independent experiments (Dunn’s multiple comparisons test, a-b p<0.0001). G) Conidiophores per field of view (5 dpi) of fungal colonies grown upon *Ecr* infection of the indicated genotypes. Mean ± SD, n=226-382 pooled from three independent experiments (Dunn’s multiple comparisons test, a-b p<0.0001. Comparison between *fer-4* and *ralf1/22/23/33*: Mann Whitney test, ** p=0.0132). All experiments were performed at least three times with similar results.

### FER and RALF mutants show post-penetration powdery mildew resistance

We next sought out to unravel cellular mechanisms underlying RALF-mediated *Ecr* susceptibility. RALF-induced calcium influx in synergid cells and pollen tubes is facilitated by FER-LRE-mediated activation of the NTA/MLO7 calcium channel (Gao *et al*., 2022). The MLO family encodes important powdery mildew susceptibility factors across families (Büschges *et al*., 1997; van Schie & Takken, 2014; Acevedo-Garcia *et al*., 2017; Nekrasov *et al*., 2017). In Arabidopsis, mutation of three MLOs is required to confer full penetration resistance (Consonni *et al*., 2006).

MLO-YFP in barley and MLO2-GFP in Arabidopsis focally accumulate at powdery mildew penetration sites (Bhat *et al*., 2005; Huebbers *et al*., 2024). We confirmed MLO2 localization to the fungal penetration site with p35S::MLO2-mCherry lines (Fig. S4A, left panel). We tested whether FER-GFP accumulates at *Ecr* penetration sites using *fer-4 pFER::FER-GFP* (Duan *et al*., 2010). Although FER-GFP signal outlined the plasma membrane, specific penetration site recruitment was not detected (Fig. S4A, right panel). The *fer-4* mutant showed strongly reduced production of conidiophores on infected tissue (Fig. 1C), but initial hyphal growth was largely unaffected (Fig. S4B). Also, epidermal cell penetration was unaffected in *fer-4* and CRISPR *ralf7x*, unlike *mlo2/6/12* (Fig. S4C). Callose deposition was not enhanced at the penetration site on *fer-4* or CRISPR *ralf7x* compared to Col-0 (Fig. S4 D, E). This suggests that mutating RALF-related components primarily affects *Ecr* reproductive success and not fungal entry or epiphytic growth. We also tested whether *Ecr* spores derived from colonies grown on *fer-4*, CRISPR *ralf7x* and *llg1-2* mutants show compromised infection potential upon re-inoculation of a Col-0 host. However, neither of the spores displayed altered pathogenicity compared to Col-0-derived spores (Fig. S4F). This indicates that, although colonies grown on the mutant plants have a reduced number of spores, their quality is not impaired.

The *mlo2/6/12* mutant displays aberrant callose deposition in trichomes with delocalized or absent accumulation (Huebbers *et al*., 2024). Likewise, *fer-4* mutants are impaired in trichome formation (Duan *et al*., 2010). Consistently, we observed less trichomes on *fer-4* but not on CRISPR *ralf7x* (Fig. S4G). Unlike *mlo2/6/12*, neither CRISPR *ralf7x* nor *fer-4* showed mis-regulated trichome callose deposition (Fig. S4H) (Huebbers *et al*., 2024). A fraction of trichomes on the CRISPR *ralf7x* mutant were enlarged in the base region which was not observed on Col-0 and *fer-4* (Fig. S4H). Further, trichomes on *fer-4* and CRISPR *ralf7x* had fewer and more branches, respectively, compared to Col-0 (Fig. S4I). Collectively, this data suggests that FER-RALF-dependent *Ecr* susceptibility and trichome development is most likely not directly linked to MLO function.

### The powdery mildew resistance phenotype of FER pathway mutants is not associated with de-regulated defense phytohormone signaling

Salicylic acid (SA) is an important defense hormone promoting resistance to biotrophic and hemibiotrophic pathogens (Peng *et al*., 2021). Ler-0 *fer-1* seedlings did not show elevated basal *PATHOGENESIS-RELATED 1* (*PR1*) expression, an SA marker gene (Kessler *et al*., 2010). Yet, the weak *fer-5* allele in Col-0 showed mildly constitutively elevated SA levels in seedlings (Duan *et al*., 2010; Engelsdorf *et al*., 2018), prompting us to test whether *fer-4’*s *Ecr* resistance phenotype may be associated with elevated SA levels. We quantified free and glycosylated SA by high performance liquid chromatography (HPLC) upon *Ecr* infection. *Ecr* infection led to mildly enhanced SA levels (Fig. 3A). However, neither *fer-4* nor CRISPR *ralf7x* showed higher SA levels before or after infection (Fig. 3A). We also measured the major catabolic SA products 2,3-DHBA and 2,5-DHBA to investigate the flux through the SA pathway (Zhang *et al*., 2013; Zhang *et al*., 2017) as well as camalexin levels associated with anti-fungal resistance (Liu *et al*., 2016; Mucha *et al*., 2019). Neither of those compounds were enhanced before or after infection in *fer-4* and CRISPR *ralf7x* (Fig. 3B, C, Fig. S5A). *PR1* expression was also not elevated in any of the mutants in control conditions. *Ecr* infection resulted in enhanced *PR1* expression. The *lrx5x* mutant showed enhanced *Ecr*-induced *PR1* expression, which, however, was not observed in the other mutants (Fig. 3D). When infected with the biotrophic oomycete pathogen *Hpa, fer-*4 and CRISPR *ralf7x* were more susceptible than Col-0 or *fer*-4/pFER::FER-GFP (Fig. S5B). This further supports the hypothesis that reduced *Ecr* sporulation observed on the tested mutants is not caused by enhanced SA signaling. Jasmonic acid (JA) is usually implicated in defense against insect herbivory and necrotrophic fungi (Bürger & Chory, 2019), but JA treatment can also inhibit colonization by the powdery mildew fungus *Erysiphe cichoracearum* (Zimmerli *et al*., 2004). FER inhibits JA signaling by destabilizing the transcription factor MYC2 (Guo *et al*., 2018). We generated a CRISPR-Cas9 *myc2* mutant in *fer-4* that abolished *MYC2* expression (Fig. S5C, D). Similar to previous reports, *fer-4* CRISPR *myc2* partially restored the *fer-4* growth phenotype (Fig. S5E) (Guo *et al*., 2018). In *fer-4*, the CRISPR *myc2* mutation showed a clear trend towards complementation of deregulated JA-responsive gene expression (Guo *et al*., 2018), a phenotype that was also observed in CRISPR *ralf7x* and *lrx5x* (Fig. 3E, F, Fig. S5D). Importantly, mutating *MYC2* did not restore *Ecr* susceptibility in *fer-4* (Fig. 3G, Fig. S5F). This suggests that the resistance phenotype of *fer-4* is independent of MYC2-mediated JA signaling.

**Figure 3:**
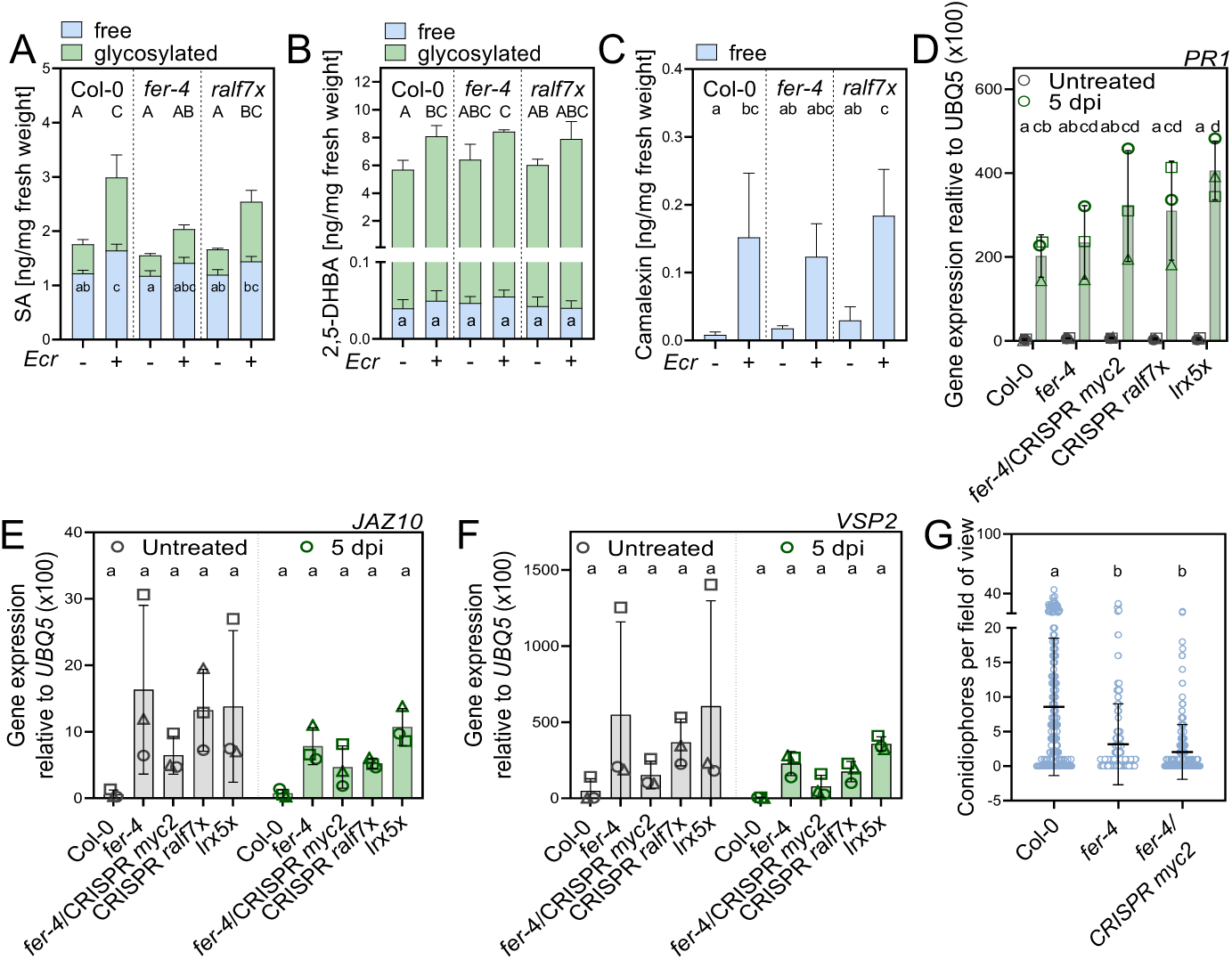
FER-RALF mediated powdery mildew susceptibility is independent of JA and SA signaling. A) Quantification of free (blue) and glycosylated (green) SA in untreated and *Ecr* infected (5 dpi) leaves of the indicated genotypes. Mean ± SD, n=3 pooled from three independent experiments (Tukey’s multiple comparisons test, free and glycosylated SA was analyzed individually, free: a-bc p=0.0423; a-c p=0.0006; ab-c p≤0.0013, glycosylated: A-C p≤0.0029; A-BC p≤0.0383). B) Quantification of free (blue) and glycosylated (green) 2,5-DHBA in untreated and *Ecr* infected (5 dpi) leaves of the indicated genotypes. Mean ± SD, n=3 pooled from three independent experiments (Tukey’s multiple comparisons test, free and glycosylated 2,5-DHBA was analyzed individually, free: not significant, glycosylated: A-C p=0.0153; A-BC p=0.0342; AB-C p=0.0345). C) Quantification of free (blue) Camalexin in untreated and *Ecr* infected (5 dpi) leaves of the indicated genotypes. Mean ± SD, n=3 pooled from three independent experiments (Tukey’s multiple comparisons test, a-bc p=0.0486; ab-c p≤0.0319; a-c p=0.0137. D) RT-qPCR of *PR1* in untreated (grey) adult leaves and upon infection with *Ecr* (5 dpi, green). Housekeeping gene *UBQ5*. Mean ± SD, n=3, data points with different symbols indicate independent biological replicates (Tukey’s multiple comparisons test, a-d p<0.0001; a-cb p≤0.0464; a-cd p≤0.0137; ab-cd p≤0.0161). E) RT-qPCR of *JAZ10* in untreated (grey) adult leaves and upon infection with *Ecr* (5 dpi, green). Housekeeping gene *UBQ5*. Mean ± SD, n=3, data points with different symbols indicate independent biological replicates (Tukey’s multiple comparisons test, not significant) F) RT-qPCR of *VSP2* in untreated (grey) adult leaves and upon infection with *Ecr* (5 dpi, green). Housekeeping gene *UBQ5*. Mean ± SD, n=3, data points with different symbols indicate independent biological replicates (Tukey’s multiple comparisons test, not significant). G) Conidiophores per field of view (5 dpi) of fungal colonies grown upon *Ecr* infection of the indicated genotypes. Mean ± SD, n=152-209 pooled from three independent experiments (Dunn’s multiple comparisons test, a-b p<0.0001). All experiments were performed three times with similar results.

### Apoplastic pH modulations affect powdery mildew infection success

To determine potential protein interaction partners of FER, we performed co-immunoprecipitation-mass spectrometry (CoIP-MS) in mature leaves of *fer-4* pFER::FER-GFP lines. We identified 68 potential interaction partners of FER-GFP with the majority being RLKs or transport-related proteins (Fig. S6A, B). We identified four CrRLK1L members (HERK1, MDS1/LET2, THE1 and LET1). In response to *Ecr* infection, we could only detect six differential FER-GFP interactors, which may be related to the early experimental time point chosen (1 dpi) (Fig. S7A, B).

Additional interactors include the plasma membrane AUTOINHIBITED H+-ATPase 1 (AHA1) and AHA3 (Fig. S6A, B). Compared to the Lti6b-GFP control, MS intensity of AHA1 was 2,7-fold enriched (Fig. 4A), confirming previously published AHA1-FER interaction in roots (Haruta *et al*., 2014; Du *et al*., 2016). We also found AHA3 to be significantly enriched with peptides only detected in FER-GFP samples (Fig. 4A, Fig. S6A, B). Perception of RALF by FER induces phosphorylation of an inhibitory site on AHA1, resulting in apoplastic alkalinization (Pearce *et al*., 2010; Haruta *et al*., 2014). Since some fungal pathogens modify the apoplastic pH for successful infection (Kesten *et al*., 2019), we hypothesized that FER-RALF-induced pH modulations may affect powdery mildew infection success. An *aha1* single mutant did not display *Ecr* colonization differences (Weis *et al*., 2013). While this might be explained by genetic redundancy, *aha1 aha2* double mutants are seedling lethal, making genetic analysis difficult (Haruta *et al*., 2010).

**Figure 4:**
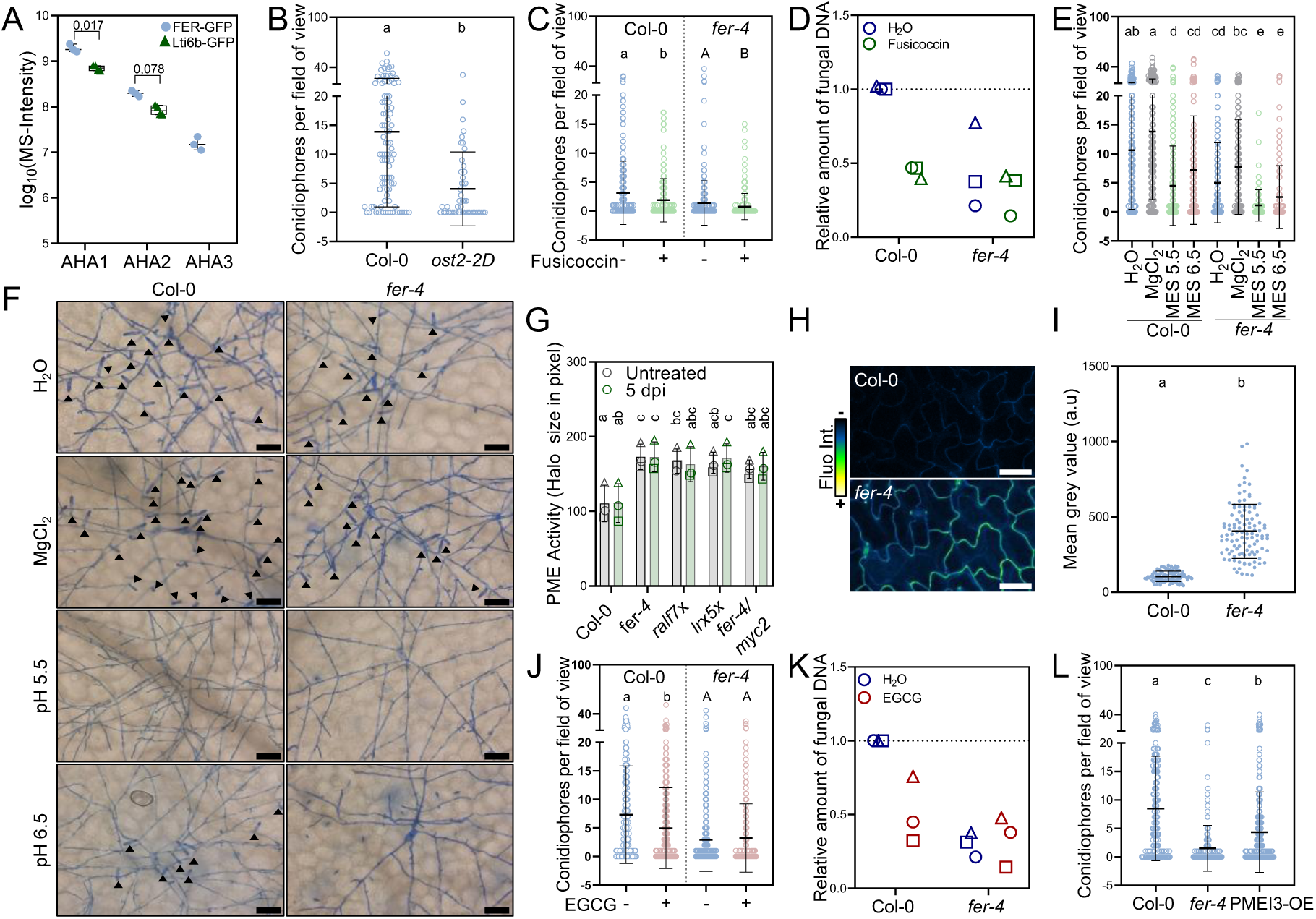
Apoplastic pH modulation and PME activity determine powdery mildew infection success. A) MS Intensities measured after Co-immunoprecipitation with untreated pFER::FER-GFP and p35S::Lti6b-GFP plants using GFP-TRAP beads. n=3 for pFER::FER-GFP, n=4 for p35S::Lti6b-GFP, data points indicate independent biological replicates. FDR values indicate statistical differences between pFER::FER-GFP and p35S::Lti6b-GFP. No AHA3 derived peptides were detected in p35S::Lti6b-GFP samples. B) Conidiophores per field of view (5 dpi) of fungal colonies grown upon *Ecr* infection of the indicated genotypes. Mean ± SD, n=49-91 pooled from two independent experiments (Mann Whitney test, a-b p<0.0001). C) Conidiophores per field of view (5 dpi) of fungal colonies grown upon *Ecr* infection of leaves pretreated with H_2_O (blue) or Fusicoccin (green). Mean ± SD, n=232-316 pooled from seven independent experiments (Mann-Whitney test, genotypes were analyzed separately, Col-0: a-b p=0.0058, *fer-4*: A-B p=0.002). D) Amount of fungal DNA normalized to plant DNA (5 dpi) upon *Ecr* infection and pretreatment with H_2_O (blue) or Fusicoccin (green). n=3, data points with different symbols indicate independent biological replicates. All data is normalized to Col-0 (H_2_O). E) Conidiophores per field of view (5 dpi) after pretreatment with H_2_O (blue), MgCl_2_ 15 mM pH 7 (grey), MES buffer 10 mM pH 5.5 (green) and MES buffer 10 mM pH 6.5 (red) before *Ecr* infection. Mean ± SD, n=126-195 pooled from three independent experiments (Dunn’s multiple comparison, a-bc, a-dc, a-d, a-e, ab-d, ab-e, bc-3 p<0.0001, ab-cd p≤0.0086, bc-d p=0.0049, d-e p≤0.034). F) Ink stained *Ecr* colonies (5 dpi) grown on Col-0 and *fer-4* plants after pretreatment with 15 mM MgCl_2_, 10 mM MES buffer (pH 5.5) and 10 mM MES buffer (pH 6.5). Black arrows indicate conidiophores. Scale bar represents 50 µm. G) PME activity as ruthenium red stained halo size in pixel in untreated leaves (grey) and upon *Ecr* infection (5 dpi, green) of the indicated genotypes. Mean ± SD, n=3, data points with different symbols indicate independent biological replicates (Tukey’s multiple comparisons test, a-c p≤0.0292, a-bc p=0.0484, ab-c p≤0.0312). H) Representative confocal images of epidermal cells of Col-0 and *fer-4* leaves stained with COS-488 labeling de-esterified pectin. Scale bar represents 50 µm. I) Quantification of COS-488 fluorescence intensity in Col-0 and *fer-4* leaves. Mean ± SD, n = 100–110 pooled from two independent experiments (Unpaired t-test, a-b p<0.0001). J) Conidiophores per field of view (5 dpi) of fungal colonies grown upon *Ecr* infection of leaves pretreated with H_2_O (blue) or EGCG (red). Mean ± SD, n=235-376 pooled from seven independent experiments (Mann-Whitney test, genotypes were analyzed separately, Col-0: a-b p<0.0001, *fer-4*: ns). K) Amount of fungal DNA normalized to plant DNA (5 dpi) upon *Ecr* infection and pretreatment with H_2_O (blue) or EGCG (red). n=3, data points with different symbols indicate independent biological replicates. All data is normalized to Col-0 (H_2_O). L) Conidiophores per field of view (5 dpi) of fungal colonies grown upon *Ecr* infection of the indicated genotypes. Mean ± SD, n=182-309 pooled from four independent experiments (Dunn’s multiple comparisons test, a-b, b-c, a-c p<0.0001). All experiments were performed at least three times with similar results.

Phosphorylation of the C-terminal penultimate threonine residue in AHA1/AHA2 (AHA1^T947^/AHA2^T948^) activates AHA proton pump activity (Falhof *et al*., 2016). We tested whether *Ecr* infection affects AHA1^T947^/AHA2^T948^ phosphorylation by using a phosphosite-specific antibody. However, pT947/pT948 phosphorylation was not altered upon *Ecr* infection (Fig. S8A, B). RALF perception by FER promotes phosphorylation of serine 899 (pS899) to inhibit AHA1/AHA2 proton pump activity (Haruta *et al*., 2014; Liu *et al*., 2025; Sun *et al*., 2025). Importantly, pS899 does not correlate with reduced pT947/pT948 and RALF23/RALF33 treatment does not affect pT947/pT948 (Haruta *et al*., 2014; Liu *et al*., 2025). Therefore, the results do not exclude a FER-dependent regulation of AHA activity correlated with altered *Ecr* infection success. To investigate a potential role of AHA1 in *Ecr* susceptibility, we tested the *ost2-2D* mutant producing a constitutive active AHA1 (Merlot *et al*., 2007). The *ost2-2D* mutant displayed enhanced *Ecr* resistance (Fig. 4B). Interestingly, the related Ler *ost2-1D* mutant displays reduced PTI responses, similar to *fer*, thus uncoupling PTI efficiency from *Ecr* infection success (Stegmann *et al*., 2017; Zhai *et al*., 2025). The fungal toxin fusicoccin induces constitutive AHA activation and promotes powdery mildew resistance in barley (Baunsgaard *et al*., 1998; Zhou *et al*., 2000). Indeed, fusicoccin treatment could also induce *Ecr* resistance and promote the *fer-4* resistance phenotype (Fig. 4C, D).

We hypothesized that alkaline pH infiltration may enhance susceptibility and/or break the *fer*-*4* resistance phenotype, whereas acidic buffers may promote resistance. We infiltrated MES buffer at pH 5.5 and pH 6.5 into the leaf apoplast prior to infection. Infiltration of a 15 mM MgCl_2_ solution (pH 7) served as an osmolarity control. Surprisingly, both infiltration of the more acidic and more alkaline buffer reduced conidiophore production in comparison to the controls, both in Col-0 and *fer*-4 (Fig. 4E, F). We next tested a more alkaline buffer and infiltrated pH 7.5 buffered HEPES into the apoplast before *Ecr* infection. In line with pH 5.5 and pH 7.5 buffered MES, pH 7.5 buffered HEPES induced strong *Ecr* resistance in both Col-0 and *fer-4* (Fig. S8C). When analyzing individual fungal colonies upon infection on Col-0 MES pH 5.5 pre-buffered leaves we observed a morphological similarity to fungal colonies on *fer*-*4* (Fig. 4F). This observation suggests the importance of apoplastic pH modulation for successful conidiophore production of *Ecr* at later time points of infection, which may be regulated through FER-RALF-dependent signaling.

Among other physiological outputs, apoplastic pH is involved in growth regulation by controlling PECTIN METHYL ESTERASE (PME) activity (Hocq *et al*., 2016; Sénéchal *et al*., 2017; Hocq *et al*., 2023). PMEs de-esterify cell wall pectins, which increases the negative charge of homogalacturonans resulting in enhanced Ca^2+^ binding and pectin cross-linking for cell wall stiffening (Willats *et al*., 2001). Importantly, demethylated pectin interacts with RALFs and LRX-RALF complexes to structurally organize the cell wall of root hairs and pollen tubes (Moussu *et al*., 2023; Schoenaers *et al*., 2024). Recently, *fer-4* mutants were shown to have constitutively increased PME activity in seedlings, which we confirmed in leaf tissue (Fig. 4G) (Biermann *et al*., 2025). Similarly, PME activity was enhanced in *lrx5x* and CRISPR *ralf7x*. *Ecr* infection, however, had no effect on PME activity in the harvested leaf samples (Fig. 4G). In accordance with previous observations in the hypocotyl (Biermann *et al*., 2025), leaf staining with the fluorescently labelled chitosan oligosaccharide probe (COS-488), which binds to de-esterified pectin oligomers (Mravec *et al.,* 2017), displayed higher signals in *fer-4* compared to Col-0 (Fig. 4H, I). Unfortunately, we could not analyze changes in the pectin methylesterification-status around *Ecr* infection sites, because of strong COS-488 staining of fungal structures (Fig. S8D). To test whether PME activity affects *Ecr* infection, we infiltrated leaves with epigallocatechin gallate (EGCG) that inhibits PMEs (Lewis *et al*., 2008). EGCG pre-treatment promoted *Ecr* resistance on Col-0 but had no effect in *fer-4* (Fig. 4J, K). To complement the inhibitor treatment, we tested the effect of PME-INHIBITOR 3 (PMEI3) overexpression on *Ecr* colonization. Overexpression of *PMEI3* inhibits RALF1 signaling in roots (Rößling *et al*., 2024). *PMEI3* overexpression resulted in reduced fungal colonization (Fig. 4L). We also tested *pme3* and *pme17* mutants because of their role in *B. cinerea* infection and FER-dependent epidermal cell lobing (Raiola *et al*., 2011; Del Corpo *et al*., 2020; Tang, W *et al*., 2022). Knockout of single *PMEs,* however, had no effect on conidiophore production (Fig. S8E). Taken together, full PME activity, which affects FER-dependent RALF signaling, is required for powdery mildew susceptibility.

### RALF-mediated powdery mildew susceptibility is partially independent of FER and FER-signaling

The reduced *Ecr* infection success upon pH buffering prompted us to resolve potential infection-induced apoplastic pH changes with cellular resolution with the ratio-metric pH sensor pUBQ10::SYP122-pHusion (Kesten *et al*., 2019). We detected apoplastic pH increases close to *Ecr* penetration sites at 2 dpi. Surprisingly, infection-related pH increases were also observed in pUBQ10::SYP122-pHusion CRISPR *fer* (Fig. 5A, B, Fig. S9A).

**Figure 5:**
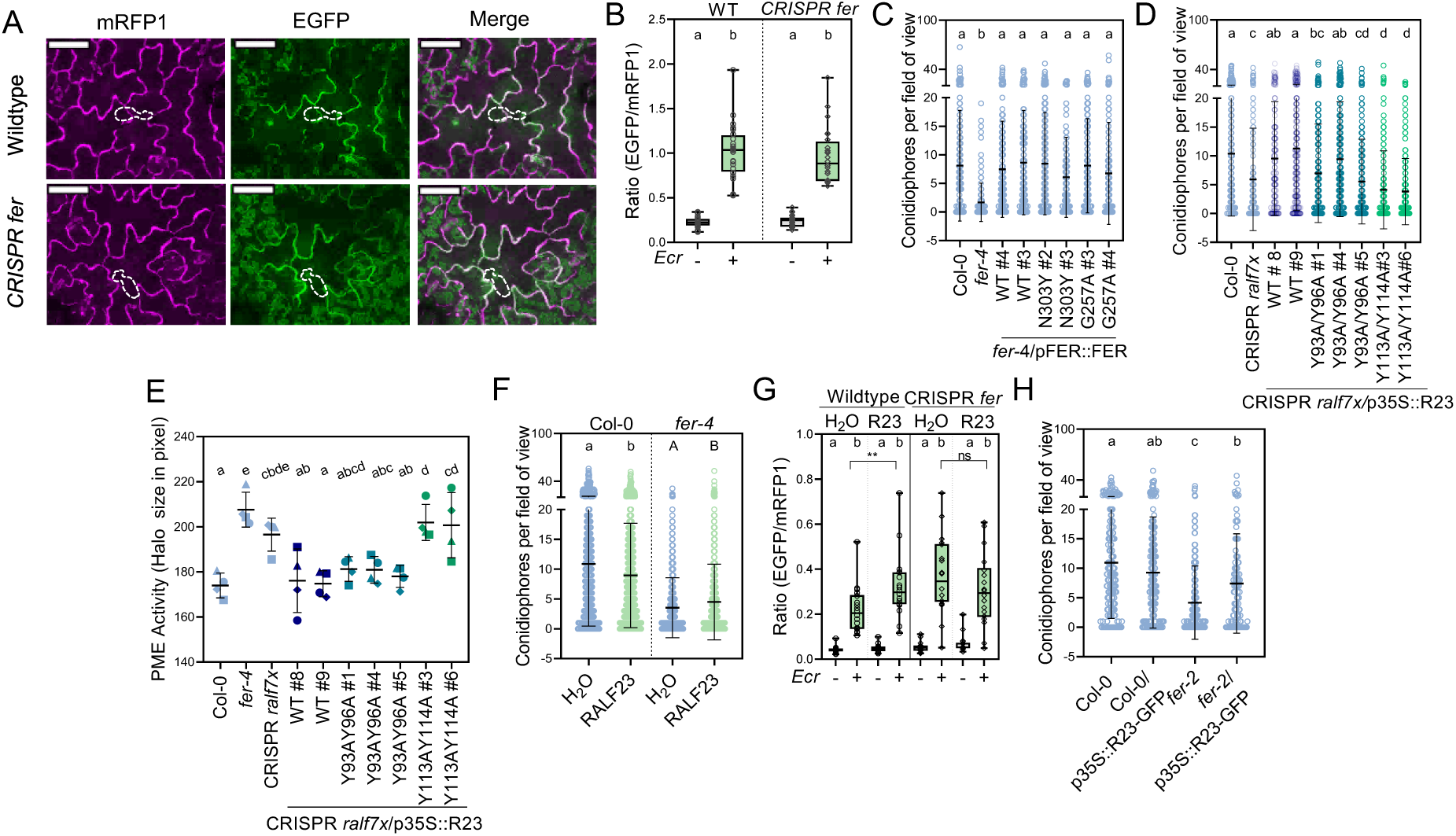
RALF-mediated powdery mildew susceptibility is partially independent of FER and FER signaling. A) Confocal images of pUBQ10::SYP122-pHusion lines in the Col-0 and CRISPR *fer* background (2 dpi). Dotted lines indicate the location of the fungal spore. Scale bar represents 50 µm. B) Quantification of the EGFP/mRFP1 ratio near the penetration site (green) and in uninfected areas (grey). Mean ± SD, n=24 pooled from three independent experiments (Mann Whitney test, a-b p<0.0001). C) Conidiophores per field of view (5 dpi) of fungal colonies grown upon *Ecr* infection of the indicated genotypes. Mean ± SD, n=131-210 pooled from four independent experiments (Dunn’s multiple comparisons test, a-b p<0.0001). D) Conidiophores per field of view (5 dpi) of fungal colonies grown upon *Ecr* infection of the indicated genotypes. Mean ± SD, n=168-326 pooled from six independent experiments (Dunn’s multiple comparisons test, a-bc p=0.0469, a-cd p<0.0001, a-d p<0.0001, ab-cc p≤0.0004, bc-dc p≤0.0019). E) PME activity normalized to Col-0. Mean ± SD, n=4, data points with different symbols indicate independent biological replicates (Tukey’s multiple comparisons test, a-e p≤0.0003, a-d p≤0.0034, a-cd p=0.0058, ab-e≤0.0012, ab-d≤0.0138, ab-cd≤0.0227, a-cbde≤0.0326). F) Conidiophores per field of view (5 dpi) after pretreatment with H_2_O (blue) or 1µM RALF23 before *Ecr* infection. Mean ± SD, n=485-970 pooled from 11 independent experiments (Mann Whitney test, statistical analysis was performed on each genotype individually, Col-0: a-b p=0.0008, *fer-4:* A-B p=0.0064). G) Quantification of the EGFP/mRFP1 ratio near the penetration site (green) and in uninfected areas (grey). 1 µM synthetic RALF peptide or H_2_O were infiltrated 2 h before infection. Mean ± SD, n=18 pooled from three independent experiments (Dunn’s multiple comparisons test a-b p≤0.0412, Mann Whitney test, **p=0.0064, ns=not significant). H) Conidiophores per field of view (5 dpi) of fungal colonies grown upon *Ecr* infection of the indicated genotypes. Mean ± SD, n=95-137 pooled from three independent experiments (Dunn’s multiple comparisons test, a-b p=0.0076, a-c p<0.0001, b-c p=0.0017). All experiments were performed at least three times with similar results.

To assess whether FER recruitment to RALF-LLG1 complexes is required for *Ecr* susceptibility, we infected *fer*-*4* lines expressing structure-guided FER mutants disrupting the LLG binding interface (Xiao *et al*., 2019). Both FER^N303Y^ and FER^G257A^ restored powdery mildew susceptibility, indicating that RALF-induced recruitment of FER to LLG1 is not essential for *Ecr* infection (Fig. 5C). We thus asked the question whether the susceptibility function of RALFs is linked to cell wall structural functions. We introduced mutations into RALF23, either disrupting the predicted LRX-binding interface (RALF23^Y113A/Y114A^), analogous to RALF4^Y83A/Y84A^ (Moussu *et al*., 2020) or the YISY motif required for LLG-FER-dependent signaling (RALF23^Y93A/Y96A^) (Xiao *et al*., 2019). We expressed both variants under control of the 35S promoter in CRISPR *ralf7x*. We tested two and three lines, respectively, with CRISPR *ralf7x* p35S::RALF23^Y93A/Y96A^ displaying similar expression levels as the WT complementation lines. CRISPR *ralf7x* p35S::RALF23^Y113A/Y114A^ lines showed weaker expression compared to the WT construct (Fig. S2F, G). RALF23^Y113A/Y114A^ could not complement the CRISPR *ralf7x* growth and enhanced *Ecr* resistance phenotype (Fig. 5D, Fig. S11). RALF23 ^Y93A/Y96A^ largely complemented the growth phenotype (Fig. S11) but only one out of three tested lines fully restored *Ecr* susceptibility (Fig. 5D). We also generated pRALF23 (pR23)::mCherry-RALF23 lines, with the mCherry tag located between the signal peptide and RALF23, carrying the same mutations. All RALF variants marked the outline of epidermal cells (Fig. S10, panel 1). Upon plasmolysis, a strong protoplastic signal for all mCherry-RALF23 variants was detected (Fig. S10, panel 2). However, mCherry-RALF23^WT^ and mCherry-RALF23^Y93A/Y96A^ displayed strong remaining signal at the cell wall, while mCherry-RALF23^Y113A/Y114A^ was fully depleted (Fig. S10, panel 1-4). This suggests loss of cell wall binding in RALF23^Y113A/Y114A^ (Fig. S10). RALF23 cell wall localization is consistent with previous reports on RALF22 and RALF4 (Moussu *et al*., 2023; Schoenaers *et al*., 2024). RALF22 secretion is impaired in *lrx1/2* (Schoenaers *et al.,* 2024). By contrast, RALF23 secretion is neither affected by disrupting LRX nor LLG1 binding motives. RALF23 and RALF23^Y93A/Y96A^, but not RALF23^Y113A/Y114A^, could rescue enhanced PME activity of CRISPR *ralf7x* (Fig. 5 E), suggesting that RALF23 interaction with LRXs is essential to inhibit PME function. Together, this data implicates that both interaction of RALF23 with LLG and LRX complexes is required to support full *Ecr* colonization with the latter playing a predominant role for RALF-regulated PME activity and powdery mildew susceptibility.

We next tested whether application of synthetic RALF23 peptide can affect powdery mildew infection success. Surprisingly, infiltration of 1 µM RALF23 into single leaves 24 h prior to *Ecr* infection had a mild resistance-inducing effect in Col-0 (Fig. 5F), while the infiltration of lower concentrations (10 nM and 100 nM) had no significant effect (Fig. S9B). Remarkably, RALF23 infiltration increased conidiophore production in the *fer-4* mutant background, suggesting that RALF23 exerts FER-independent effects (Fig. 5F). Interestingly, infiltration with RALF23 promoted alkalinization near the infection site in pUBQ10::SYP122-pHusion, but not in CRISPR *fer* (Fig. 5G). Surprisingly, water infiltration (control) increased *Ecr*-induced alkalinization in CRISPR *fer.* This raises the question whether FER is additionally involved in buffering *Ecr*-induced pH changes upon apoplast infiltration. Also, RALF23 infiltration promoted GFP fluorescence in pUBQ::SYP122-pHusion in a FER-dependent manner at early time points post treatment. Buffering abolished the recorded pH shifts (Fig. S9C), suggesting that it effectively compromises RALF23 pH changes. We continued testing the effect of RALF23-GFP overexpression on *Ecr* infection outcome. Similar to exogenous RALF23 treatment (Fig. 5F), RALF23-GFP overexpression displayed a trend towards enhanced *Ecr* resistance in Col-0 (Fig. 5H). In *fer-2,* RALF23-GFP overexpression promoted susceptibility (Fig. 5H) and displayed a clear trend to reduce the mutant’s enhanced PME activity (Fig. S9D). This observation is in line with previous findings that RALF23 treatment significantly reduces PME activity in *fer-4*, but not in Col-0 (Biermann *et al*., 2025). *RALF23-GFP* showed overexpression in both Col-0 and *fer-2* (Fig. S9D).

Collectively, our data suggests that RALF-mediated powdery mildew susceptibility is at least partially FER-independent. Moreover, it indicates that likely appropriate spatio-temporal availability of RALFs affect *Ecr* reproductive success in a combination of partially FER-dependent effects on alkalinization and FER-independent responses, potentially related to RALF-regulated PME activity.

## Discussion

This work provides first molecular insights into the mechanism of FER-mediated powdery mildew susceptibility. We revealed that FER’s RALF peptide ligands are critical for the completion of *Ecr*’s asexual life cycle upon host colonization.

RALFs are perceived by LLG-FER/CrRLK1L complexes to induce downstream signaling and bind to cell wall localized LRX proteins and demethylated pectin for cell wall organization. Our data suggests that powdery mildew sporulation requires functional apoplastic pH homeostasis and modifications of the cell wall pectin methyl-esterification status, both processes in which FER-RALF and/or LRX-RALF modules fulfil central regulatory functions. Earlier work revealed that barley powdery mildew infection is accompanied by pH shifts and the induction of apoplastic alkalinization at early time points post inoculation (2h) (Felle *et al*., 2004). Importantly though, mildew-induced pH shifts were dynamic, with fluctuations between alkaline and more acidic conditions during the recorded time period (50 h) (Felle *et al*., 2004). In light of our observations, disrupting apoplastic pH homeostasis by buffering, RALF23 peptide treatments or RALF23 overexpression, may circumvent dynamic endogenous regulatory circuits and inhibit *Ecr* host colonization. A challenge for the future will be to reveal spatio-temporal pH fluctuations and RALF secretion during powdery mildew infection.

In addition, the pectin methyl-esterification status contributes to powdery mildew pathogenesis. Both EGCG treatment and PMEI3 overexpression promoted *Ecr* resistance. Notably, RALF-signaling outputs and the methyl-esterification status of the cell wall are tightly interconnected. PMEI3 is strongly pH dependent with the highest activity at acidic conditions (Xu *et al*., 2022) RALF-induced apoplastic alkalinization may inhibit PMEI3 which increases PME activity for cell wall remodeling. RALF22 and RALF4 bind to LRXs to affect pectin compaction in the cell wall (Moussu *et al*., 2023; Schoenaers *et al*., 2024). In return, PME activity modulates RALF1/RALF23 signaling and the formation of RALF1-pectin molecular condensates required for FER-LLG1-dependent responses (Liu *et al*., 2024; Rößling *et al*., 2024; Biermann *et al*., 2025). Consistently, EGCG treatment enhanced powdery mildew resistance in Col-0 but had no additive effect in *fer-4*. Considering that *fer-4* is strongly compromised in RALF perception, our data indicates that PME inhibition through EGCG treatment affects the same FER-dependent pathway. We propose that EGCG treatment and PMEI3 overexpression block sustained *Ecr* growth by reducing the availability of demethylated pectin for RALF signaling through FER. Fusicoccin and buffer treatment could also induce resistance in *fer-4*, suggesting additional effects that do not solely rely on pH-dependent PME regulation. Our work revealed that *fer*, CRISPR *ralf7x* and *lrx5x* display elevated steady-state PME activity, in accordance with previous findings (Biermann *et al*., 2025). Moreover, PME activity is primarily regulated by RALF binding to LRXs. This raises the question whether disruption of the FER-RALF-LRX pathway activates feedback mechanisms to promote PME activity in an effort to re-establish RALF cell wall binding and signaling. Interestingly, RALF23 treatment and overexpression in *fer* can reduce PME activity (Biermann *et al*., 2025) and enhance powdery mildew susceptibility. This indicates FER independent effects of RALF23 on cell wall organization and powdery mildew infection. FER is dispensable for RALF22-induced root hair tip alkalinization, calcium influx and cell wall modification (Schoenaers *et al*., 2024) and RALF23 affects cell wall pectin status in a partially FER-independent and LRX-dependent manner (Biermann *et al*., 2025). Our SYP122-pHusion results suggest that *Ecr* induces apoplastic pH shifts independent of FER, raising the question of receptor redundancy or RALF-independent pH modulations, possibly through PTI (Zhai *et al*., 2025; Wang *et al*., 2026) or, alternatively, actively triggered by the fungus. Interestingly though, RALF23-induced enhancement of *Ecr*-triggered apoplastic pH modulation is FER dependent. This is in line with our findings that RALF23 interaction with FER-LLG1 supports full colonization. However, RALF23 binding to LRXs is important for cell wall binding and essential for *Ecr* infection success. This suggests a predominant role for RALF23 structural function and/or effects on cell wall chemistry to support *Ecr* colonization. Yet, the interplay between RALF structural and signaling function remains largely unknown (Schade *et al*., 2025). RALFs bind to LRX with high affinity at acidic pH, while LLGs require alkaline conditions for optimal RALF recognition (Moussu *et al*., 2020). It is possible that dynamic apoplastic pH fluctuations induced by *Ecr* infection releases RALFs from LRXs to trigger CrRLK1L-dependent signaling that further affects its reproductive success. Our data also raises the question whether additional FER-related receptors are required for RALF perception to establish full *Ecr* susceptibility. FER can interact with related RALF-binding CrRLK1Ls, including HERK1 and ANJ, to form higher order complexes (Galindo-Trigo *et al*., 2020; Zhong *et al*., 2022; Lan *et al*., 2023). This is supported by our CoIP-MS data according to which FER interacted with several members of the CrRLK1L family, namely HERK1, MDS1, THE1 and LET1.

The fungal haustorium is surrounded by the extrahaustorial matrix (EHMx) and the extrahaustorial membrane (EHM), a plant derived membrane that is continuous with the plasma membrane but has distinctive properties (Hückelhoven & Panstruga, 2011; Polonio *et al*., 2021). At the site of invasion the EHMx is sealed with densely packed cell wall depositions forming the haustorial neck, rendering the EHMx a separate apoplastic compartment (Micali *et al*., 2011). Nutrients taken up from the EHMx should derive from the invaded plant cell and require prior export. Plant cell sugar uptake is facilitated by SUGAR TRANSPORT PROTEINs (STP), which are associated with plant microbe interactions (Bezrutczyk *et al*., 2018). In Arabidopsis, STP8-GFP, upon overexpression, localizes to the EHM and increases susceptibility to powdery mildew (Liu *et al*., 2021) and transport-compromised STP13 alleles are associated with resistance to powdery mildew in wheat and barley (Milne *et al*., 2019). STPs are H^+^ symporters, and STP8 has a pH optimum at 5.5 for cellular import (Liu *et al*., 2021). It is possible that *Ecr* infection induces RALF-CrRLK1L-dependent changes to the apoplastic pH that reprograms transporter topology to favor haustorial sugar uptake over plant cell import. In accordance, *fer* mutants show altered sucrose uptake and starch accumulation (Yang *et al*., 2015; Yeats *et al*., 2016). Supporting our hypothesis, we identified several transporter proteins as FER interactors in our CoIP-MS experiment, including STP1, involved in resistance against bacterial infection (Yamada *et al*., 2016). We did not identify many differential FER-GFP interactors upon *Ecr* infection, likely due to the early experimental time point chosen (1 dpi).

We propose a model in which RALF-mediated apoplastic pH-modulation and pectin de-methylesterification are a prerequisite for sustained powdery mildew host colonization (Figure 6). In accordance, endogenous pectin modification affects powdery mildew infection outcome. Mutants in the pectate lyase *POWDERY MILDEW RESISTANT 6 (PMR6)* and the pectin acetyltransferase *PMR5* are more resistant to powdery mildew without altered fungal penetration or de-regulated SA/JA signaling (Vogel *et al*., 2002; Vogel *et al*., 2004). Our data shows that RALF pathway mutants display a similar phenotype. Moreover, the *fer-4* mutant has reduced cell wall cellulose content (Yeats *et al*., 2016) and dysfunctional apoplastic pH modulation which could interfere with fungal nutrient uptake. This supports a previously coined hypothesis of altered fungal nutrition as a consequence of cell wall defects (Vogel *et al*., 2002; Vogel *et al*., 2004).

**Figure 6:**
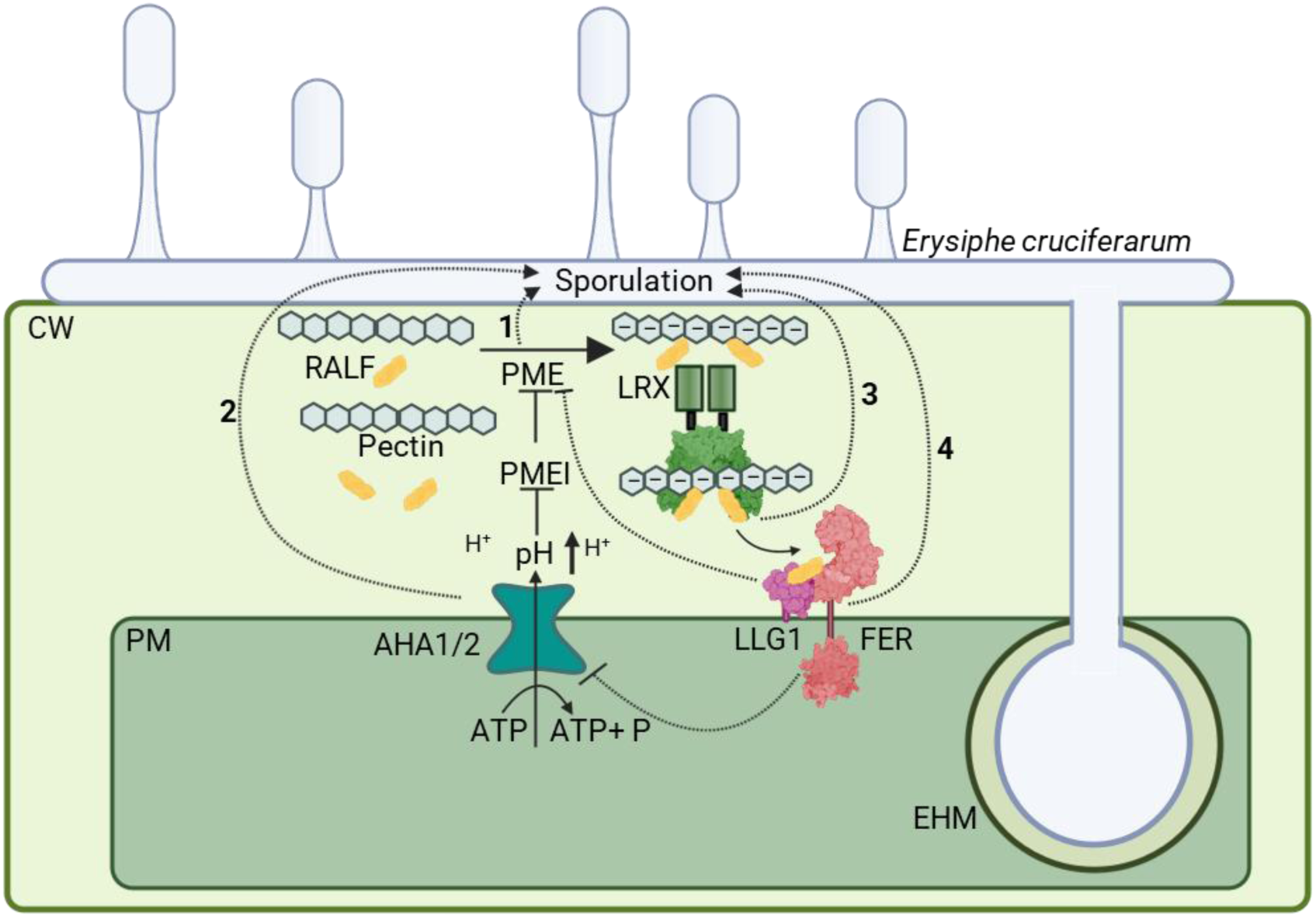
Model of RALF-FER-dependent effects on powdery mildew sporulation. RALF peptides bind to negatively charged pectins and LRXs in the cell wall, as well as the FER-LLG1 complex at the plasma membrane. RALF binding to FER-LLG1 results in inhibition of AHA1/2 activity and apoplastic pH changes. The apoplastic pH modulates PMEI/PME activity and with this likely the methyl-esterification status of cell wall pectins. Both PME activity (1) and apoplastic pH modulation (2) affect powdery mildew sporulation. RALF-peptides appear to affect sporulation in a FER-independent (3) and FER-dependent (4) manner, potentially through regulating cell wall structure and the apoplastic pH. CW: cell wall; PM: Plasma membrane, EHM: Extrahaustorial membrane. Created with BioRender.com.

Other plant pathogens produce RALFs to establish host colonization, including *F. oxysporum* and root knot nematodes (Masachis *et al*., 2016; Zhang *et al*., 2020). Interestingly, these RALF peptide mimics lack conserved sequence elements required for LRX binding, but contain a modified YISY motif required for LLG-CrRLK1L-dependent signaling (Pearce *et al*., 2010). This suggests that fungal RALFs support virulence mainly through CrRLK1L-induced signaling and less through cell wall binding. None of the analyzed powdery mildew genomes contained RALF peptide mimics, suggesting that *Ecr* likely relies on the combination of endogenous RALF signaling and cell wall remodeling to support sustained host interaction. Our work thus highlights a new mode of RALF function, combining signaling and structural roles for the control of plant microbe interactions.

## Materials and Methods

### Powdery mildew propagation

*Erysiphe cruciferarum* was propagated weekly by transferring spores to 4 – 5-week-old uninfected Col-0 plants. Spores were used for further inoculation 3 – 4 weeks after propagation. Propagation and experiments were performed in an environmentally controlled growth cabinet (22 °C, 70 % relative humidity, 12 h photoperiod).

### Molecular cloning

To generate p35S::RALF23 and p35::MLO2-mCherry lines gene sequences were cloned from gDNA using primers listed in Table S3 and assembled into a GoldenGate-modified pCB302 binary vector for plant expression. The p35S::RALF23^Y93A/Y96A^ and p35S::RALF23^Y113A/Y114A^ constructs were obtained by site-directed mutagenesis using primers listed in Table S3 and likewise assembled into a GoldenGate-modified pCB302 binary vector. For pR23::mCherry-RALF23 constructs, pR23 was cloned using primers listed in Table S3. Gene fragments for RALF23^WT^, RALF23^Y93A/Y96A^ and RALF23^Y113A/Y11A^ with an N-terminal mCherry-tag located between the signal peptide and the mature RALF23 were obtained by gene synthesis (Twist Bioscience, USA). pR23 and the synthesized gene fragments were assembled into a GoldenGate-modified pCB302 binary vector for plant expression. To generate CRISPR-Cas9 mutants the software tool chopchop (https://chopchop.cbu.uib.no/) was used to design target sites (1 to 2 per gene of interest). All target sequences are listed in Table S4. Individual guide RNAs with gene specific target sites were obtained by gene synthesis (Twist Bioscience, USA). Higher order gRNA stacks were cloned into pICSL4723OD with FastRed-pRPS5::Cas9 (Castel *et al*., 2019). All generated plant expression constructs were transformed into *Agrobacterium tumefaciens* strain GV3101 before floral dip transformation.

### Plant material and growth conditions

Plant lines used in this study are listed in Table S2. All plants were grown with two plants per pot in environmentally controlled conditions (21 °C, 55 – 65% relative humidity, 8 h photoperiod). 5 – 6-week-old plants were used for *Ecr* infection.

### Conidiophore counting and fungal penetration assay

Conidiophore production was analyzed 5 days after *Ecr* infection. Leaves were destained using EtOH:Acetic Acid (6:1) and subsequently stained using ink (ink: 25 % acetic acid, 9:1). Fungal structures were visualized using a Zeiss AXIO imager Z1.m microscope. Fungal penetration success was similarly analyzed and quantified at 2 dpi. A penetration attempt was scored as successful when the spore formed a penetration tube and first hyphal structures.

### Quantification of fungal DNA by qPCR

For quantification of fungal DNA, two infected plants per genotype were harvested at 5 dpi before DNA extraction with standard protocols. RT-qPCR experiments were performed using Maxima SYBR green mix (Thermo Fisher Scientific, USA) on an AriaMx Real-Time PCR system (Agilent Technologies, USA). The amount of fungal DNA was calculated relative to plant DNA using primers for the Arabidopsis small RubisCO subunit (AtRbcS) and *Ecr* tubulin (Table S3).

### Reinoculation experiments

5 weeks old Col-0, *fer-4*, *llg1-2* and CRISPR *ralf7x* plants were highly infected with powdery mildew. 11 days post infection spores from these plants were used to infect 5 weeks old Col-0.

### Hyaloperonospora arabidopsidis (Hpa) infection and quantification

For *Hpa* infection assays, plants were vernalized for 2 – 3 days in the dark at 4 °C and grown in environmentally controlled conditions (22 °C, 60% relative humidity, 10 h photoperiod). 14-day-old seedlings were spray-inoculated with a spore suspension of *Hpa* isolate Noco2 (5 × 104 spores/mL) (Asai *et al*., 2015). After inoculation, plants were kept in trays covered with sealed lids to maintain high humidity (18 °C, near-saturation humidity). At 7 days post inoculation, infected aerial parts were harvested in water and vortexed to release spores. Spores were counted using a hemocytometer and normalized to leaf fresh weight.

### Infiltration experiments

Individual leaves were infiltrated with synthetic RALF23 (Sequence: ATRRYISYGALRRNTIPCSRRGASYYNCRRGAQANPYSRGCSAITRCRRS) 24 h prior to infection. Fusicoccin (10 µM), EGCG (50 µM) and MES buffers were infiltrated 2 h before *Ecr* infection.

### Gene expression analysis

For RNA extraction, TRIzol reagent (Roche, Switzerland) and purification with Direct-zol™ RNA Miniprep Plus kit (Zymo Research, Germany) with on column DNAse I digestion was used. RNA was reverse transcribed with random hexamer primer and Revert Aid reverse transcriptase (Thermo Fisher Scientific, USA). RT-qPCR experiments were performed using Maxima SYBR green mix (Thermo Fisher Scientific, USA) on an AriaMx Real-Time PCR system (Agilent Technologies, USA). Gene expression levels were normalized to *UBIQUITIN5* (*UBQ5*). All primers are listed in Table S3.

### PME activity assay

PME activity assays were performed as previously described (Bethke *et al*., 2014). In brief, plant material of untreated and infected plants was harvested 5 dpi and ground in liquid nitrogen. Plant powder was incubated with 500 µL protein extraction buffer (50 mM Tris-HCl pH 7.5, 150 mM NaCl, 10% glycerol, 5 mM dithiothreitol, 1% protease inhibitor cocktail, 2 mM Na2MoO4, 2.5 mM NaF, 1.5 mM activated Na3VO4, 1mM phenylmethanesulfonyl fluoride, 0.5% IGEPAL) for 30 min at 8°C before centrifugation at 16 000 g for 30 min. The supernatant was transferred into a fresh 1,5 mL tube and protein concentration was determined by Bradford assay. The protein concentration was adjusted to 3 µg/µL. To assess PME activity, the 15 µL liquid was loaded into holes in an agarose gel plate containing esterified pectin (1.2 % (w/v) agarose, 0.1 % (w/v) pectin from apple, 12 mM citric acid, 50 mM Na_2_HPO_4_ pH 7) before incubation at 37°C for 16 h. Afterwards, plates were briefly washed with water and stained with 0.05 % ruthenium red for 30 min. Residual dye was washed off with water and images of the plates were captured by scanning. Darker stained areas correlating with PME activity were quantified using Fiji ImageJ software (Schindelin *et al*., 2012).

### AHA-phosphorylation analysis

Infected plant material was ground in liquid nitrogen. Protein extraction buffer (3 % [w/v] SDS, 30 mM TRIS-HCl pH 8.0, 10 mM EDTA. 10 mM NaF, 30 % [w/v] Sucrose, 0.012% [w/v] Coomassie Brilliant Blue, 15 % [v/v] 2-Mercaptoethanol) and extracted proteins were immediately used for SDS- PAGE/western blot. Proteins were detected using AHA/pThr947 antibodies (Hayashi *et al*., 2010). Band intensities were quantified using Fiji ImageJ software.

### Quantification of salicylic acid, dihydroxybenzoic acids and camalexin

SA, DHBAs and camalexin were extracted as described (Nawrath & Métraux, 1999) with minor modifications. Ground fine leaf powder was spiked with 250 ng *ortho*-anisic acid (oANI) as internal standard and sequentially extracted with 70% and 90% methanol for 1 hour at 65°C. Combined supernatants were concentrated, macromolecules precipitated with 5 % trichloroacetic acid and supernatants partitioned two times against cyclohexane / ethyl acetate (1:1) to obtain free SA, DHBAs and camalexin. The aqueous phase was acidified with 8 M HCl (1:1) and incubated for 1 hour at 80°C. Hydrolyzed extracts were spiked with oANI and partitioned as above to obtain glycosylated SA and DHBAs. Organic phases of free and glycosylated fractions were concentrated and dissolved in 90% 25 mM KH_2_PO_4_, pH 2.6 / 10% acetonitrile (ACN). HPLC analysis was performed on an Agilent 1260 Infinity II system with fluorescence detection. SA, DHBAs, camalexin and oANI were separated on an Agilent InfinityLab Poroshell 120 SB-C18 column (3 x 100 mm, 2.7 µm) at 40°C, using a solvent gradient of (A) 25mM KH_2_PO_4_, pH 2.6 and (B) ACN:H_2_O (99:1) at 1 mL/min flow rate (all steps in v/v): 0 to 1 min: 10% B; 1 to 6 min: 10 – 25 %B; 6 to 8.5 min: 25 – 80 %B; 8.5 to 10.5 min: 80% B; 10.5 to 11 min: 80 – 10% B; 11 to 14 min 10% B. SA (Ex 305 nm, Em 407 nm), DHBAs (Ex 320 nm, Em 450 nm), camalexin (Ex 318 nm, Em 370 nm) and oANI (Ex 305 nm, Em 365 nm) were calibrated against a standard dilution series of all metabolites of interest in the range of 1 to 100 ng.

### Histochemical staining of callose deposition in rosette leaves and trichomes

Three rosette leaves per 35-d-old Arabidopsis plant were collected and destained in 70% ethanol (v/v). Afterwards, leaves were incubated over-night in aniline blue solution (0.01% Aniline Blue (m/v) in 150 mM K_2_HPO_4_ buffer) and mounted on a microscopy slide. Micrographs were captured by a BZ-9000 microscope (Keyence, Osaka, Japan) using UV illumination. Images were optimized by Adobe Photoshop 2024 in an identical manner for all micrographs. Trichome and trichome branch number were scored by counting.

### Callose quantification at the fungal penetration site

Leaves were harvested and destained in EtOH:Acetic Acid (6:1). Next, leaves were incubated for 10 min in 0.067 mM K_2_HPO_4_. Subsequently, leaves were transferred into staining solution with 0.05% methyl blue in 0.067 M K_2_HPO_4_ and incubated overnight. Fungal structures were stained before analysis with 0.05 % propidium iodide (PI). Methyl blues fluorescence was detected between 480 - 530 nm with a laser at 458 nm for excitation. PI fluorescence was excited with a 510 nm laser and detected at 600 – 620 nm. Z-Projections were created and mean grey values were measured using Fiji (Schindelin *et al*., 2012).

### COS-488 synthesis, imaging and quantification

COS-488 was synthesized as described before (Mravec *et al*., 2017). Chitosan oligosaccharides (Carbosynth OC09272) were dissolved in 100 mM NaCH_3_CO_2_ (pH 4.9) to 1 mg/mL. AlexFluor 488 hydroxylamine was prepared in DMSO (10 mg/mL).16 µL were added to 500 µL COS solution and incubated at 37°C in dark for two days under gentle shaking. Staining of leaf epidermis cells was performed by harvesting leaf disks from mature Arabidopsis plants. COS-488 was infiltrated at 1:1000 dilution in ½ MS + 1% Sucrose pH 5.8 under mild vacuum. Afterwards, leaf disks were incubated in COS-488 solution for 2 h and washed with ½ MS + 1 % Sucrose pH 5.8 three times under mild vacuum. Samples were mounted and z-stacks acquired using a Leica SP8 confocal microscope equipped with HC PL APO 40x water objective, argon laser and HyD detectors. COS-488 was excited at 488 nm. Emission was detected between 490-560 nm. Z-Projections were created and mean grey values were measured using Fiji (Schindelin *et al*., 2012).

### Confocal microscopy

Confocal laser-scanning microscopy was performed using a Leica TCS SP5 (Leica, Germany) microscope (with Leica Application Suite X 3.7.4.23463). All images were taken with a 20 x water immersion objective. To visualize GFP fluorescence an argon laser with HyD detector was used (emission at 488 nm, detection 500-550 nm). mCherry fluorophore was excited with a DPSS laser (emission 561 nm, detection 600-640 nm). and a detection window of 600 - 640 nm. Changes in the apoplastic pH around fungal infection sites were analyzed using pUBQ10::SYP122-pHusion expressing plants. GFP and RFP were excited with a 488 nm and a 561 nm laser, respectively. Signals were detected at 500 – 545 nm (GFP) and 650 – 695 nm (RFP). Z-stacks with identical settings were taken at the penetration site and in unpenetrated areas. The GFP/RFP ratios were determined using Fiji Image J.

### CoIP-MS

Sample preparation and mass spectrometric data acquisition for all CoIP samples was carried out as previously described (Scheinost *et al*., 2026). Protein identification and quantification were performed with MaxQuant software, using parameters previously described (Scheinost *et al*., 2026) and the following exceptions: Proteins were identified by searching MS2 spectra against the TAIR protein database for *Arabidopsis thaliana* (downloaded March 2022, 48231 protein entries). Phosphorylation of serine/threonine/tyrosine, oxidation of methionine and acetylation at the protein N-terminus were specified as variable modifications. The “match-between-run” functionality was enabled (matching time window 0.7 min, alignment time window 20 min). For protein quantification the MaxQuant output table proteinGroups.txt and Label-Free Quantification (LFQ) (Cox et al., 2014) intensities were used. Only proteins detected in at least two biological replicates in at least one experimental condition were further processed. Missing values were imputed by a protein-specific constant value, which was defined as the lowest detected protein-specific LFQ-value over all samples divided by two, using the data visualization platform omicsViewer (DOI: 10.1101/2022.03.10.483845). Additionally, a maximal imputed LFQ value was defined as 15% quantile of the protein distribution from the dataset. The volcano plots (Figure S6A, S7A) were generated using RStudio. Interactions were marked as significant if they passed the threshold of FDR < 0.05 and fold change > 2. The FER-GFP infected with *Ecr* and untreated experiments were performed in biological triplicates, while all other experiments were performed in biological quadruplets.

### Identification of putative powdery mildew RALF-likes

To identify putative RALF-like peptides in the predicted proteins and genome assemblies of selected phytopathogenic fungi, we used the hmmsearch program from the hmmer suite v3.3.2 (www.hmmer.org) with default parameters. The hidden markov model profile (.hmm) for the RALF motif (PF05498.15) was obtained from the PFAM website (https://ftp.ebi.ac.uk/pub/databases/Pfam/current_release/Pfam-A.hmm.gz), access data 19.11.2023). RALF HMM profile was used to search predicted protein sequences from the selected fungal species (see Supplementary Table S1) using the hmmsearch function. Hits with E-values below 1e^-05^ were retained. To perform the hmmsearch analysis on the genome assemblies, we used a custom Python script to split the contigs into 300 bp fragments with a sliding window of 50bp. All six open reading frames of these split sequences were translated and stop codons were removed to generate a multi-FASTA file suitable for be processed with hmmsearch. Search hits with E-values < 1e-05 were retained. Accession number of genome assemblies used in this analysis are reported in Supplementary Table S1.

### Quantification and statistical analysis

GraphPad Prism (Version 8.0.1) was used to perform statistical analysis. Detailed descriptions of the sample size, p-values and statistical methods are indicated in the respective figure legends.

## Acknowledgements

We thank Stefanie Ranf for providing GoldenGate vectors for molecular cloning and Mark Youles (TSL Norwich) and Laurence Tomlinson (TSL Norwich) for providing the pICSL4723OD vector for CRISPR-Cas9 cloning. We thank Hannah König for supporting the collection of micrographs to assess callose depositions in rosette leaves and trichomes. The authors thank Verena Breitner and Franziska Hackbarth for excellent technical assistance and maintenance of mass spectrometers at the BayBioMS. Confocal imaging was performed at the Center for Advanced Light Microscopy (CALM) of the TUM School of Life Sciences. This work was funded by the Deutsche Forschungsgemeinschaft (DFG) grants STE2448/4-1 (HL and MS), TRR356 TP B09 (HL, SS and MS), TRR356 TP B01 (JG), TRR356 TP B08 (MKRL and AM), SFB1101 TP A08 (JG), individual research grant 451218338 (MKRL), EN 1071/3-1 (TE) and PA861/20-1 (JH). The Q Exactive HFX mass spectrometer was funded in part by the DFG grant INST 95/1435-1 FUGG (CL). The work was further supported by the French government through the France 2030 investment plan managed by the National Research Agency (ANR), as part of the Initiative of Excellence of Université Côte d’Azur under reference number ANR-15-IDEX-01 (ABD), the Technical University of Munich (MS, HL, DB, JG, SS, MM, RH) and Ulm University (MS).

## Author contributions

Conceptualization: MS; Investigation: HL, SS, JH, KM, GH, DB, AM, XZ, YC, MM, ABD, MS; Funding acquisition: MS, RH, ABD, MKRL, JG, TE, TK, MM, CL; Project administration: MS; Supervision: MS, HL, RH, ABD, MKRL, JG, TE, CL; Writing – original draft: HL, MS; Writing – review & editing: all authors.

## Data availability

The mass spectrometric raw files as well as the MaxQuant output files have been deposited to the ProteomeXchange Consortium via the PRIDE partner repository and can be accessed using the identifier PXD059737 (https://proteomecentral.proteomexchange.org/cgi/GetDataset?ID=PXD059737). Reviewer account username, password: lumM0fo5d8wq.

---

**Figure S1:**
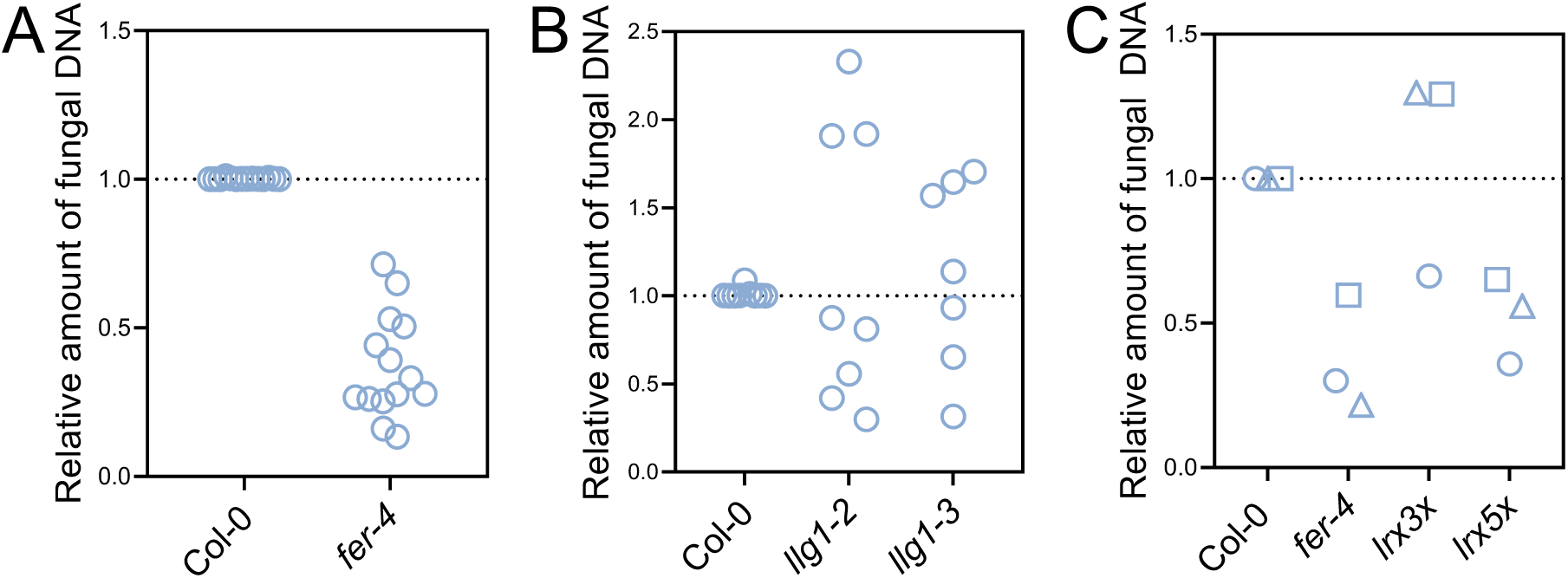
Quantification of fungal growth on mutants of RALF binding proteins. A-C) Amount of fungal DNA normalized to plant DNA (5 dpi) upon *Ecr* infection of the indicated genotypes. Mean ± SD, data points indicate independent biological replicates. A) n=14, B) n=7–9, C) n=3. C) Data points with different symbols indicate independent biological replicates. All experiments were done at least three times with similar results.

**Figure S2:**
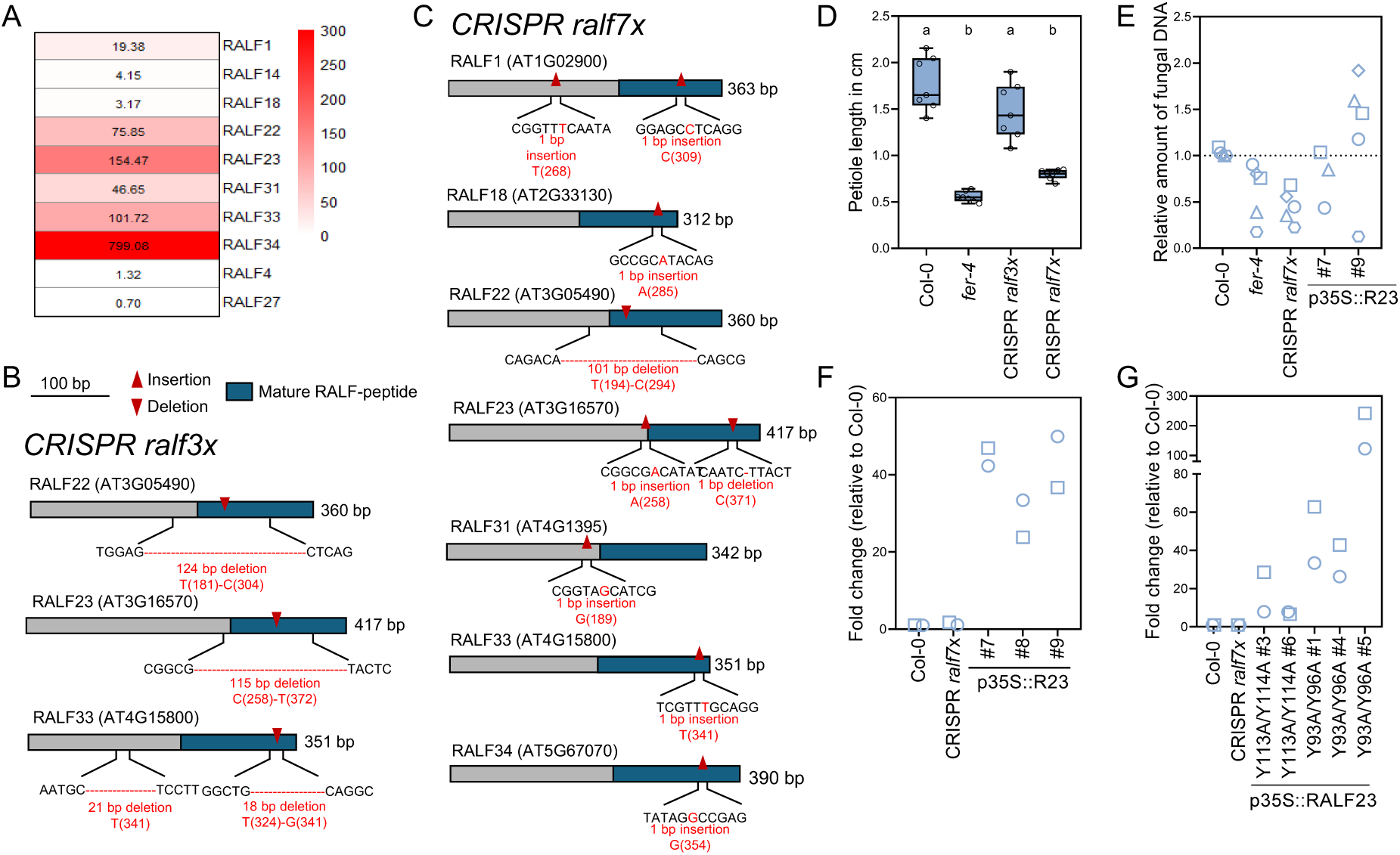
Characterization of CRISPR *ralf* mutants. A) Expression levels of the indicated *RALF* genes in the vegetative rosette. Data was obtained using ePlant browser (https://bar.utoronto.ca/eplant/) and is based on (Schmid *et al*., 2005). B) Characterization of CRISPR *ralf3x*. Schematic diagram of *RALF22, RALF23* and *RALF33* gene structure and the CRISPR-Cas9-mediated mutation pattern detected by DNA sequencing. The mature RALF peptide is indicated in blue. C) Characterization of CRISPR *ralf7x*. Schematic diagram of *RALF1, RALF18, RALF22, RALF23*, *RALF31, RALF33* and *RALF34* gene structure and the CRISPR-Cas9-mediated mutation pattern detected by DNA sequencing. The mature RALF peptide is indicated in blue. D) Quantification of the petiole length of 4-week-old Col-0, *fer-4*, CRISPR *ralf3x* and CRISPR *ralf7x* plants. Each data point represents the average length of petioles on a fully grown rosette. Mean ± SD, n=7 pooled from two independent experiments (Tukey’s multiple comparisons test, a-b p<0.0001). E) Amount of fungal DNA normalized to plant DNA (5 dpi) upon *Ecr* infection of the indicated genotypes. Data points with different symbols indicate independent biological replicates n=3-5, F-G) RT-qPCR of *RALF23* in adult leaves of the indicated plant lines. Housekeeping gene *UBQ5,* n=2. Data points with different symbols indicate independent biological replicates. D-E) were performed at least two times with identical results. F–G) were performed twice with identical results.

**Figure S3:**
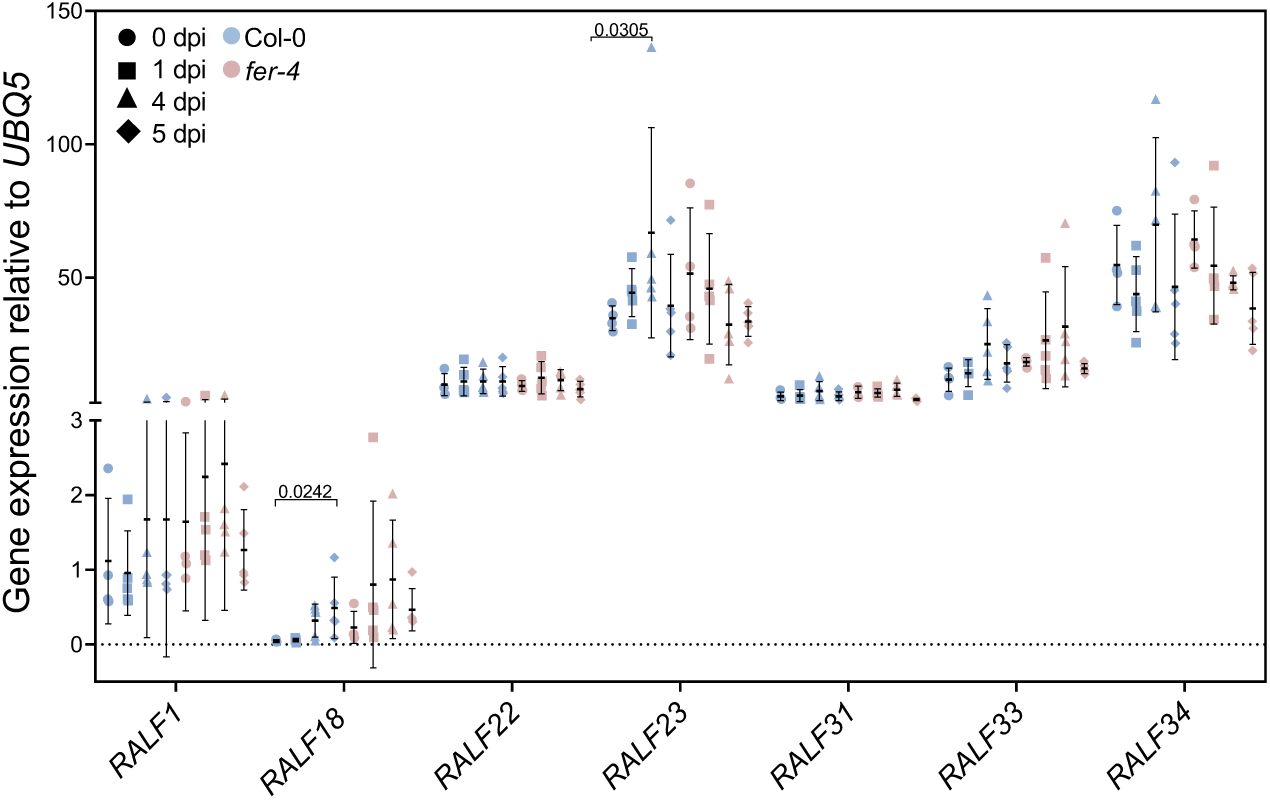
Regulation of RALF peptides during powdery mildew infection. RT-qPCR of the indicated *RALF* genes in Col-0 (blue) and *fer-4* (red) at 0 dpi (circle), 1 dpi (square), 4 dpi (triangle), 5 dpi (diamond). Housekeeping gene *UBQ5*. Mean ± SD, n=4-5, data points indicate independent biological replicates. (Dunn’s multiple comparison test against 0 dpi within the respective genotype, p-values < 0.05 are indicated in the graph, all other comparisons were not significant.

**Figure S4:**
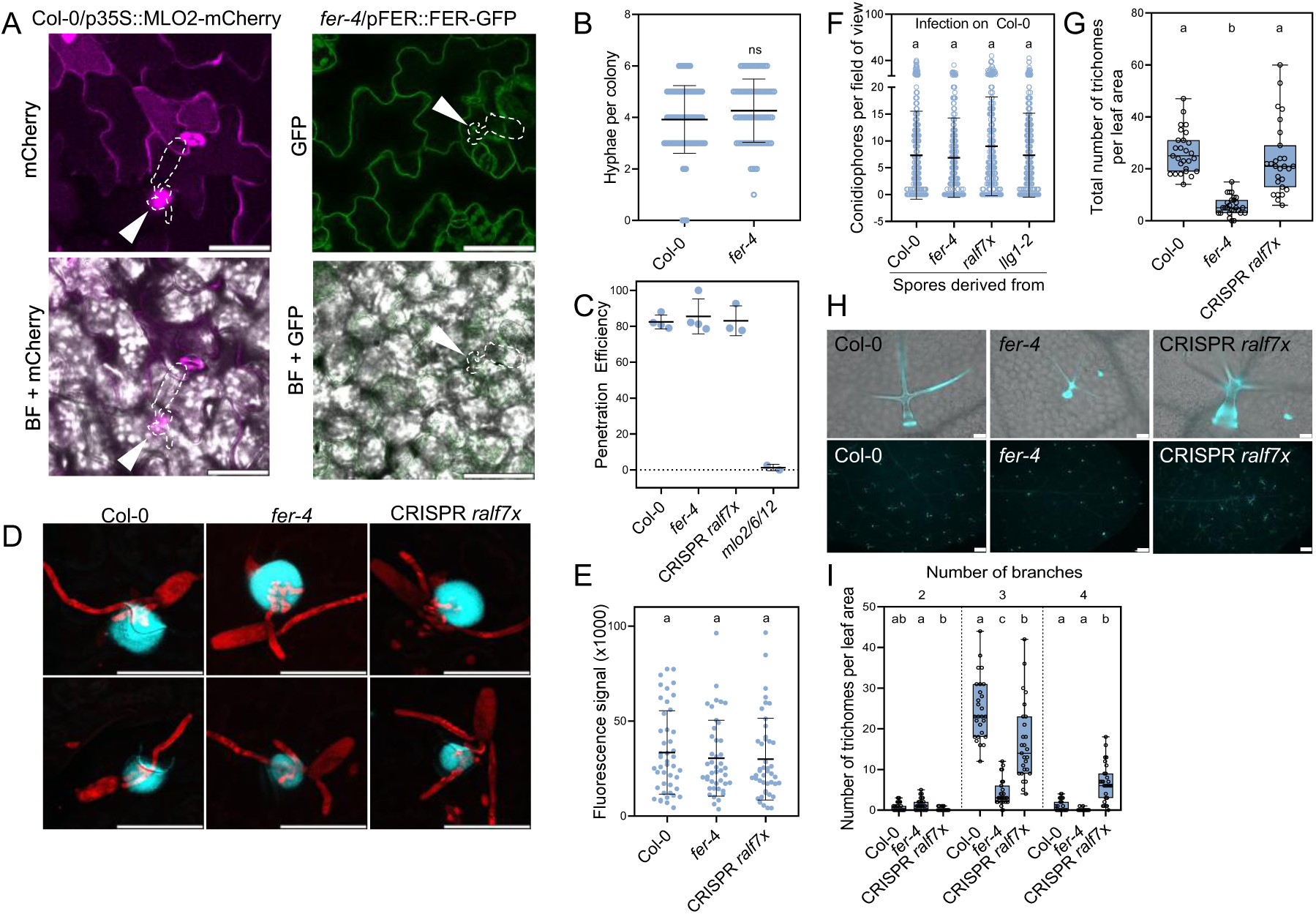
FER- and RALF-mediated powdery mildew susceptibility is most likely not directly linked to MLO. A) Confocal microscopy images of fungal spores penetrating p35S::MLO2-mCherry and pFER::FER-GFP lines (2 dpi). Dotted lines indicate fungal spores, white arrow marks the penetration site. Scale bar represents 50 µm. B) Quantification of hyphal branching (2 dpi). Mean ± SD, n=77–98 pooled from two independent experiments (Mann Whitney test, ns: not significant). C) Penetration efficiency of *Ecr* on the indicated genotypes (calculated as the percentage of successful penetrations from all counted interactions, 2 dpi). Mean ± SD, n=2–4. D) Confocal microscopy images of fungal spores at 1 dpi. Callose depositions were stained using methyl blue (turquoise). Fungal structures were stained with propidium iodide (red). Scale bar represents 50 µm. E) Quantification of methyl blue fluorescence around the fungal penetration site, indicating the amount of callose deposition. Mean ± SD, n=43-45 pooled from three independent experiments (Dunn’s multiple comparisons test). F) Conidiophores per field of view (5 dpi) of fungal colonies grown on Col-0 plants. Spores from the indicated genotypes were used for infection. Mean ± SD, n=218-336 pooled from three independent experiments (Dunn’s multiple comparisons test). G) Quantification of trichome numbers per leaf area on the indicated genotypes. Mean ± SD, n=27 pooled from three independent experiments (Dunn’s multiple comparisons test, a-b p<0.0001). H) Microscopic images of aniline blue-stained trichomes of Col-0, *fer-4* and CRISPR *ralf7x*. Scale bar in the upper row represents 50 µm, scale bar in the lower panel represents 500 µm. I) Quantification of trichome branching on the indicated genotypes. Mean ± SD, n=27 pooled from three independent experiments (Dunn’s multiple comparisons test, statistical analysis was performed on each category individually, 2 branches: a-b p=0.0002; 3 branches: a-b p=0.0259; a-c, b-c p<0.0001; 4 branches: a-b p<0.0001). All experiments were performed three times with similar results, except B), which was performed twice with identical results.

**Figure S5:**
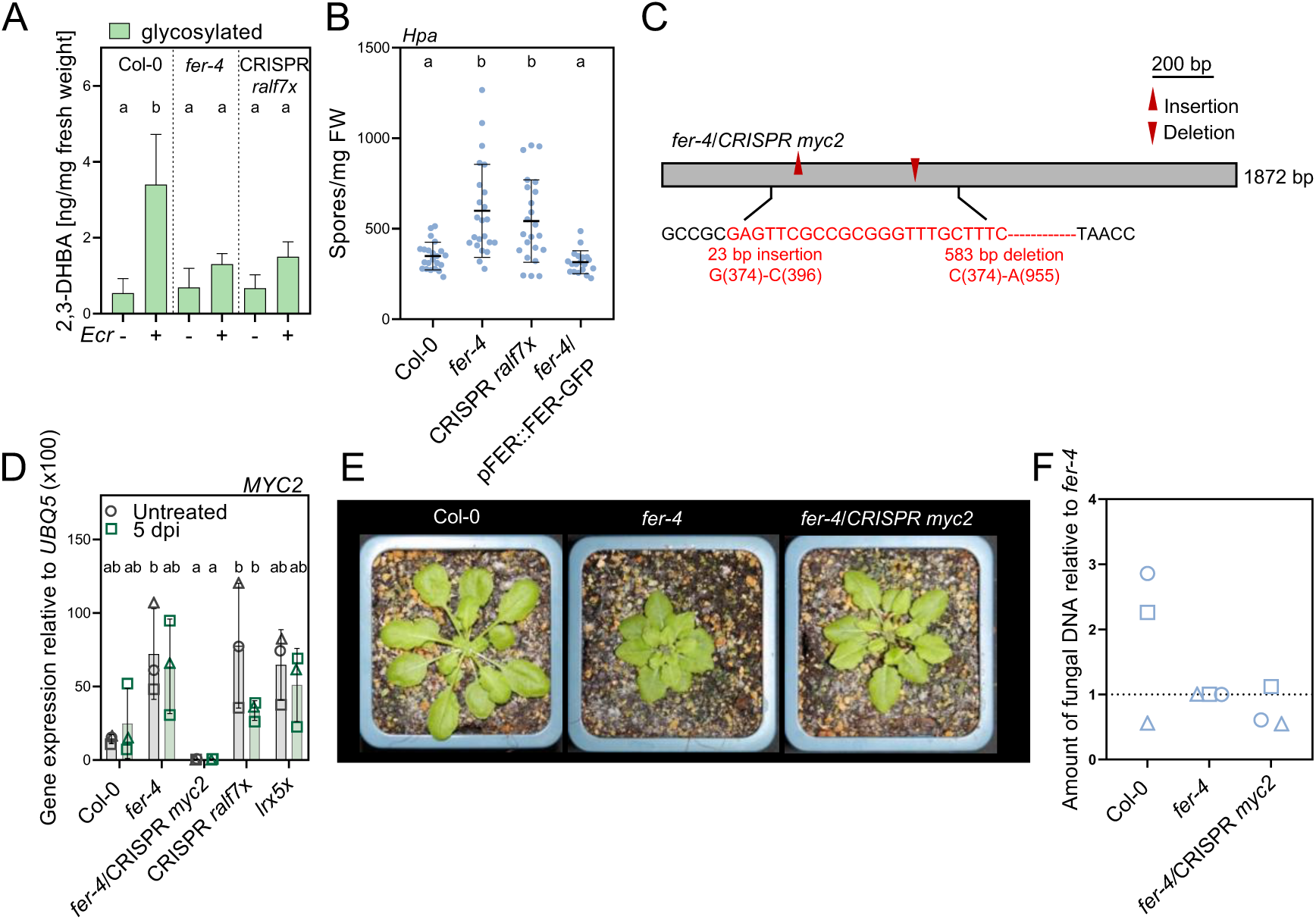
**Characterization of *fer-4* CRISPR *myc2.*** A) Quantification of glycosylated (green) 2,3-DHBA in untreated and *Ecr* infected (5 dpi) leaves of the indicated genotypes. Mean ± SD, n=3 pooled from three independent experiments (Tukey’s multiple comparisons test, glycosylated: a-b p≤0.0324). B) Fungal spores/mg fresh weight (FW) of *Hyaloperonospora arabidopsidis* (*Hpa*) grown on the indicted genotypes. Mean ± SD, n=19-23 pooled from three independent experiments (Dunn’s multiple comparison test, a-b p≤0.0129). All experiments were repeated at least three times with similar results. C) Characterization of *fer-4* CRISPR *myc2*. Schematic diagram of *MYC2* gene structure and the CRISPR-Cas9-mediated mutation pattern detected by DNA sequencing. D) RT-qPCR of *MYC2* in untreated (grey) adult leaves and upon infection with *Ecr* (5 dpi, green). Housekeeping gene *UBQ5*. Mean ± SD, n=3, data points with different symbols indicate independent biological replicates (Tukey’s multiple comparisons test, a-b p≤0.0358). E) Images of 4-week-old Col-0, *fer-4* and *fer-4* CRISPR *myc2* plants. F) Amount of fungal DNA normalized to plant DNA (5 dpi) upon *Ecr* infection of the indicated genotypes, n=3. Data points with different symbols indicate independent biological replicates. A, B, D and F were performed three times with similar results.

**Figure S6:**
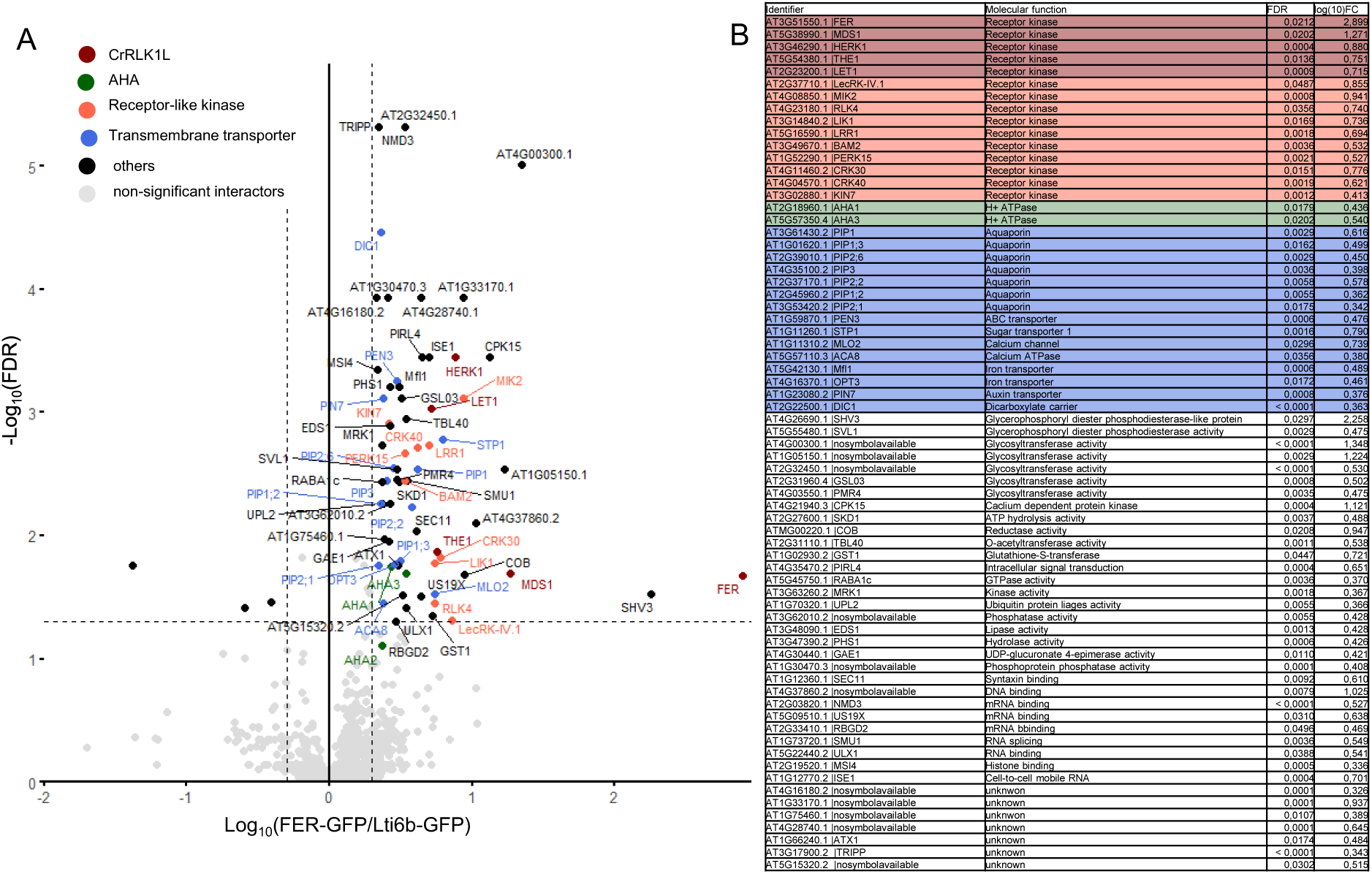
FER interactors detected by CoIP-MS. A) Volcano plot of detected FER-GFP interactors. Significantly enriched proteins (FDR=0.05 and fold change (FER-GFP/Lti6b-GFP) >2) are marked with colored circles according to the legend. Non-significant proteins are colored grey. B) List of significant FER-GFP interactors compared to Lti6b-GFP control. Data was obtained from biological triplicates for FER-GFP and biological quadruplets for Lti6b-GFP.

**Figure S7:**
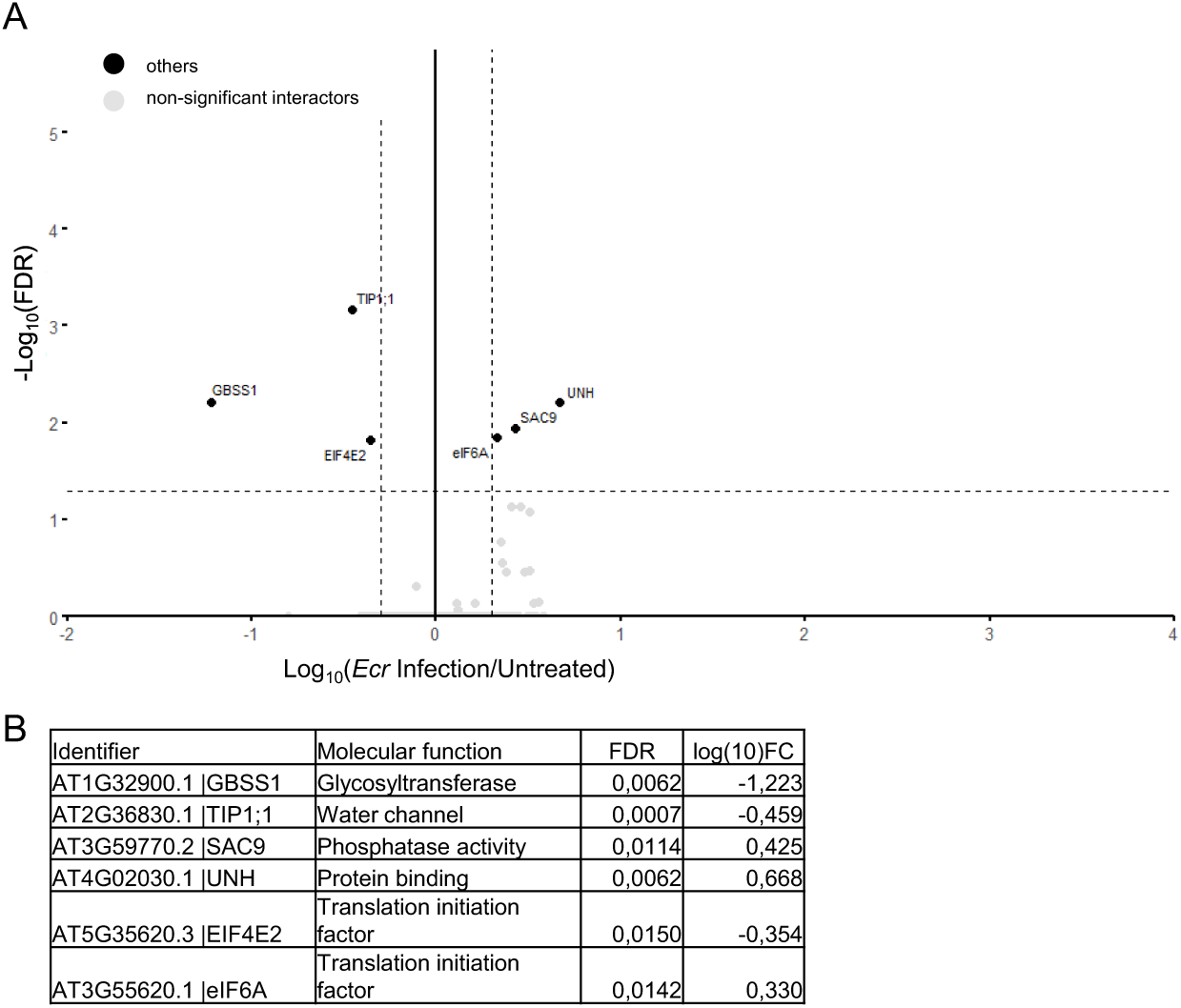
Powdery mildew dependent FER interactors detected by CoIP-MS. A) Volcano plot of detected FER-GFP interactors after *Ecr* infection (1 dpi). Significantly enriched proteins (FDR=0.05 and fold change (*Ecr* infection (1dpi)/Untreated) >2) are marked in black. Non-significant proteins are colored grey. B) List of significant FER-GFP interactors at 1 dpi with *Ecr*, compared to the untreated FER-GFP control. Data was obtained from biological triplicates for each treatment.

**Figure S8:**
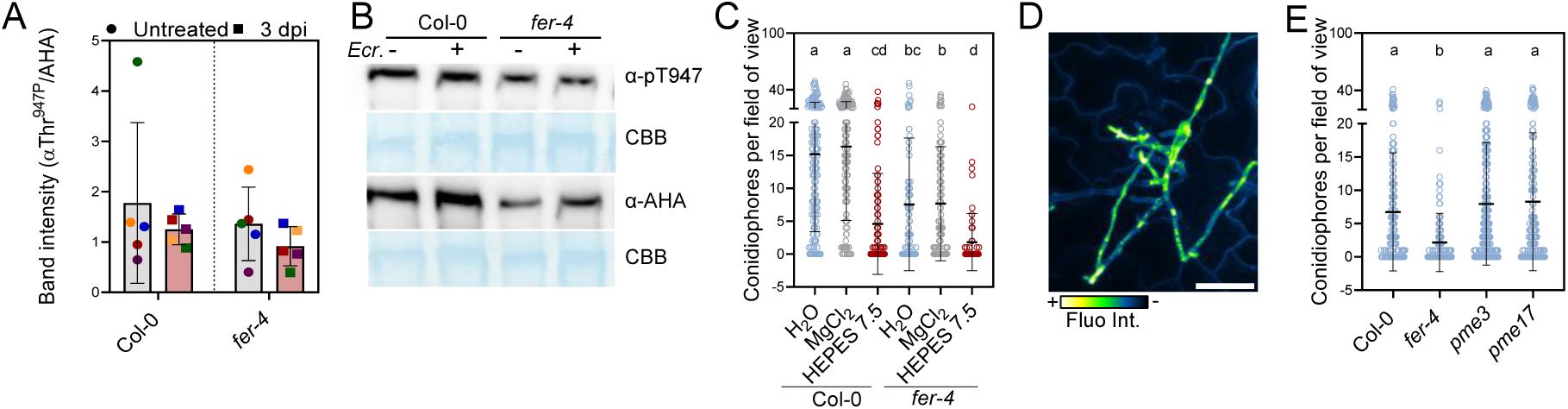
Role of apoplastic pH and PMEs for *Ecr* infection success. A) Quantification of the band intensities detected in western blot probed with α-AHA and α-Thr^947P^ antibodies. Band intensities were measured using Image J software. Data points indicate individual experiments. Data points from the same experiment are marked with the same color. B) AHA phosphorylation upon infection with *Ecr* (3 dpi). Western blots were probed with α-AHA and α-Thr^947P^ antibodies. CBB = Coomassie brilliant blue. C) Conidiophores per field of view (5 dpi) after pretreatment with H_2_O (blue), MgCl_2_ 10 mM pH 7 (grey), HEPES buffer 10 mM pH 7.5 (red) before *Ecr* infection. Mean ± SD, n=53-153 pooled from three independent experiments (Dunn’s multiple comparison, a-b, a-dc, a-bc, a-cd, a-d, b-d p<0.0001, b-cd p=0.0245). D) Confocal images of fungal structures stained with COS-488. Scale bar represents 50 µm E) Conidiophores per field of view (5 dpi) of fungal colonies grown upon *Ecr* infection of the indicated genotypes. Mean ± SD, n=161-338 pooled from three independent experiments (Dunn’s multiple comparisons test, a-b p<0.0001). All experiments were repeated at least three times with similar results.

**Figure S9:**
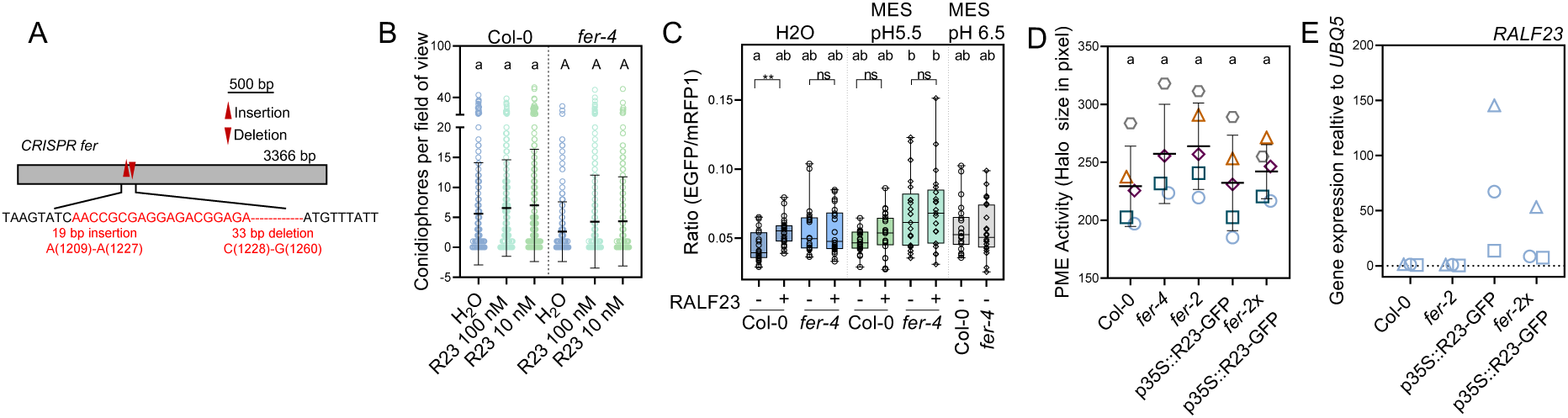
Analysis of FER-independent RALF23 functions during *Ecr* infection. A) Characterization of pUBQ10::SYP122-pHusion CRISPR *fer-4*. Schematic diagram of *FERONIA* gene structure and the CRISPR-Cas9-mediated mutation pattern detected by DNA sequencing. B) Conidiophores per field of view (5 dpi) after pretreatment with H_2_O (blue), 100 nM RALF23 (turquoise), or 10 nM RALF23 (green). Mean ± SD, n=210-259 pooled from four independent experiments (Dunn’s multiple comparisons test, genotypes were analyzed separately. C) Quantification of the EGFP/mRFP1 3 h post infiltration of 1 µM RALF23 in H_2_O or MES pH 5.5. Mean ± SD, n=21-24 pooled from three independent experiments (Dunn’s multiple comparisons test a-b≤0.009, Mann Whitney test, **p=0.0017, ns=not significant). D) PME activity normalized to Col-0. Mean ± SD, n=4-5, data points with different symbols indicate independent biological replicates (Tukey’s multiple comparisons test, not significant). (E) RT-qPCR of *RALF23* in untreated adult leaves of the indicated plant lines. Housekeeping gene *UBQ5*. n=3, data points with different symbols indicate independent biological replicates. B-E) All experiments were performed at least three times with similar results.

**Figure S10:**
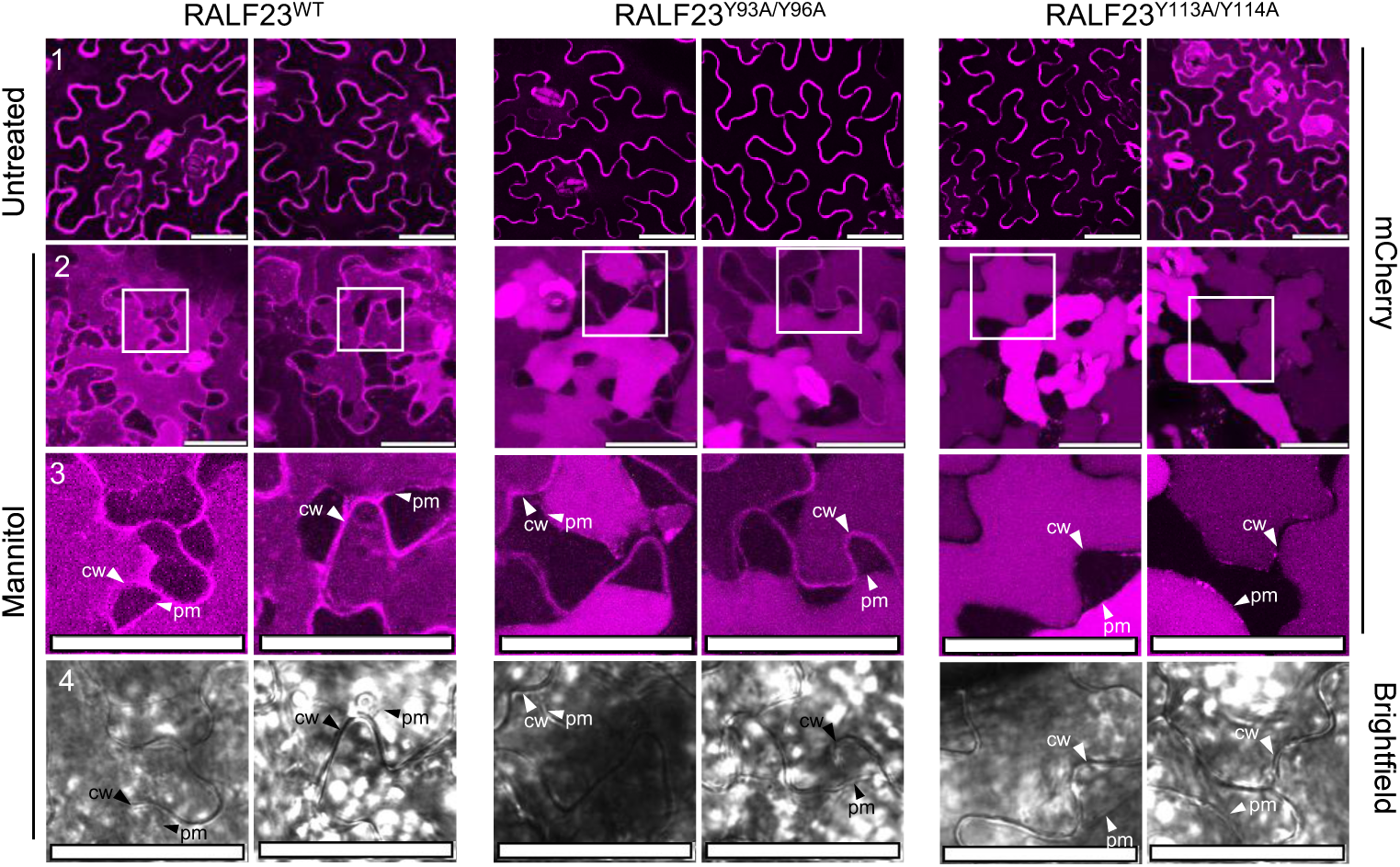
Cell wall localization of mCherry-RALF23 peptide variants. A) Representative z-stack images of epidermal cell expressing different variants of pR23::mCherry-RALF23. Panel 1: mCherry-RALF23 localization in untreated epidermal cells, panel 2-4: Epidermal cells after mannitol treatment (0.5 M), panel 3 and 4 show the image section marked in panel 2. Scale bar represents 50 µm. cw: Cell wall, pm: Plasma membrane.

**Figure S11:**
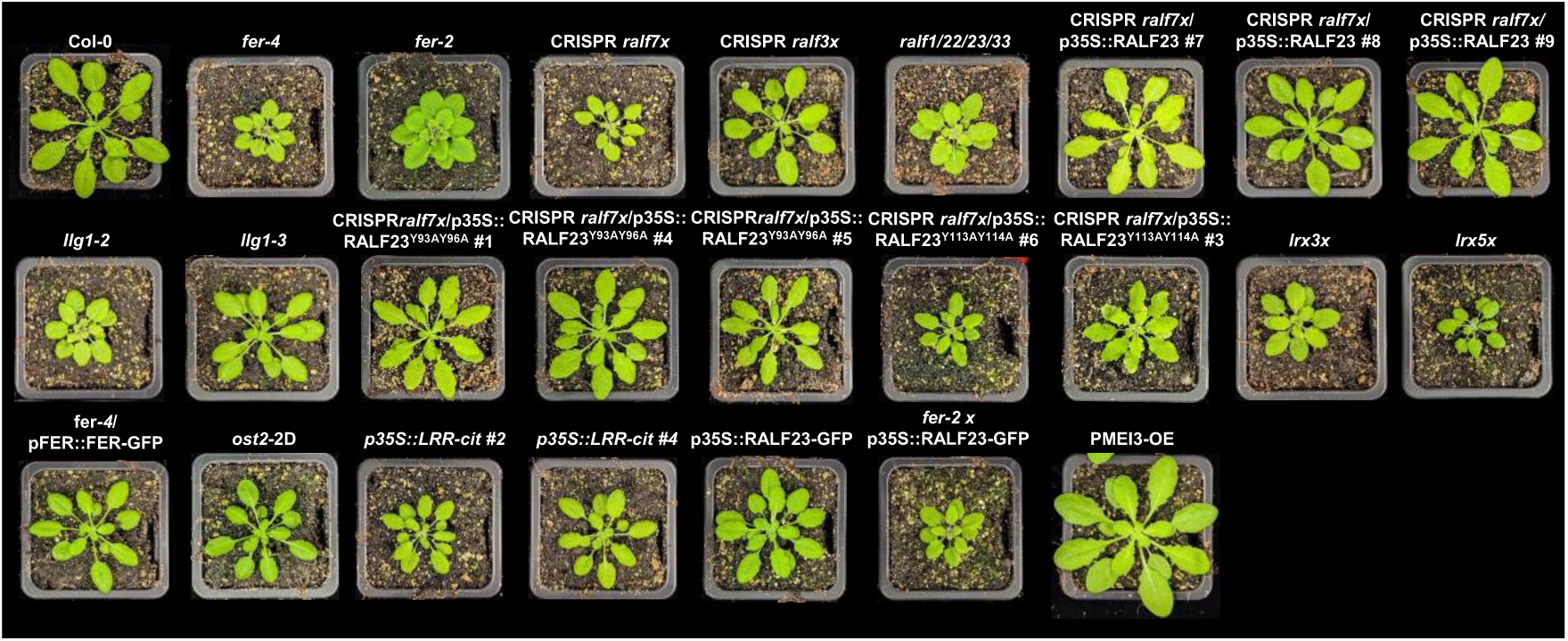
Images of plant lines used in this study. Representative images of 5-week-old plants used in this study.

**Table S1:** Identification of RALF-like peptides from phytopathogenic fungi species reported to be able to infect Arabidopsis.

| Species | Significant hits in<br>predicted proteins<br>(E-values < 10e<sup>-05</sup>) | Significant hits in<br>translated genome<br>assembly (E-values<br>< 10e<sup>-05</sup>) | GenBank assembly<br>accession number | Reference |
| --- | --- | --- | --- | --- |
| *Fusarium oxysporum*<br>NRRL 32931 | 1 | 2 | GCA_000271745.2 | (Delulio *et al.*,<br>2018) |
| *Pyrenophora tritici-*<br>*repentis* Pt-1C-BFP | 1 | 1 | GCA_000149985.1 | (Manning *et al.*,<br>2013) |
| *Golovinomyces*<br>*cichoracearum*<sup>a</sup> UMSG1 | 0 | 0 | GCA_003611235.1 | (Wu *et al.*, 2018) |
| *Erysiphe*<br>*neolycopersicum*<sup>a</sup><br>UMSG2 | 0 | 0 | GCA_003610855 | (Wu *et al.*, 2018) |
| *Erysiphe necator*<br>EnFRAME01 | 0 | 0 | GCA_024703715.1 | (Zaccaron *et al.*,<br>2023) |
| *Blumeria graminis* f.sp.<br>*tritici* CHE_96224 | 0 | 0 | GCA_900519115.1 | (Müller *et al.*,<br>2019) |
| *Podosphaera xanthii*<br>YZU573 | 0 | 0 | GCA_028751805.1 | (Xu & Chen,<br>2023) |

**Table S2:** Plant lines used in this study.

| Plant line | Genetic Background | Reference |
| --- | --- | --- |
| *fer-4* | Col-0 | (Duan *et al.*, 2010) |
| *fer-2* | Col-0 | (Deslauriers & Larsen, 2010) |
| pFER::FER-GFP | *fer-4* | (Duan *et al.*, 2010) |
| *llg1-2* | Col-0 | (Li *et al.*, 2015) |
| *llg1-3* | Col-0 | (Shen *et al.*, 2017) |
| *lrx1/2/3* | Col-0 | (Draeger *et al.*, 2015) |
| *lrx1/2/3/4/5* | Col-0 | (Herger *et al.*, 2020) |
| p35S::LRR4-citrine | Col-0 | (Fabrice *et al.*, 2018) |
| *ralf1/22/23/33* | Col-0 | (Lan *et al.*, 2023) |
| CRISPR *ralf3x* | Col-0 | Generated in this study |
| CRISPR *ralf7x* | Col-0 | Generated in this study |
| p35S::RALF23 | CRISPR *ralf7x* | Generated in this study |
| p35S::RALF23<sup>Y93A/Y96A</sup> | CRISPR *ralf7x* | Generated in this study |
| p35S::RALF23<sup>Y113A/Y114A</sup> | CRISPR *ralf7x* | Generated in this study |
| pR23::mCherry-RALF23 | CRISPR *ralf7x* | Generated in this study |
| pR23::mCherry-RALF23<sup>Y93A/Y96A</sup> | CRISPR *ralf7x* | Generated in this study |
| pR23::mCherry-RALF23<sup>Y113A/Y114A</sup> | CRISPR *ralf7x* | Generated in this study |
| *mlo2/6/12* | Col-0 | (Consonni *et al.*, 2006) |
| CRISPR *myc2* | *fer-4* | Generated in this study |
| PMEI3-OE | Col-0 | (Peaucelle *et al.*, 2008) |
| *ost2-2D* | Col-0 | (Merlot *et al.*, 2007) |
| *pme3* | Col-0 | (Guénin *et al.*, 2011) |
| *pme17* | Col-0 | (Del Corpo *et al.*, 2020) |
| pUBQ10::SYP122-pHusion | Col-0 | (Kesten *et al.*, 2019) |
| pUBQ10::SYP122-pHusion/CRISPR *fer* | pUBQ10::SYP122-pHusion | Generated in this study |
| pFER::FER-GFP | *fer-4* | (Chakravorty *et al.*, 2018) |
| pFER::FER<sup>K565R</sup>-GFP | *fer-4* | (Chakravorty *et al.*, 2018) |
| p35S::RALF23-GFP | Col-0 | (Dobón *et al.*, 2015) |
| *fer-2* x p35S::RALF23-GFP | *fer-2* | (Stegmann *et al.*, 2017) |
| pFER::FER<sup>WT</sup>-GFP | *fer-4* | (Xiao *et al.*, 2019) |
| pFER::FER<sup>N303Y</sup>-GFP | *fer-4* | (Xiao *et al.*, 2019) |
| pFER::FER<sup>G257A</sup>-GFP | *fer-4* | (Xiao *et al.*, 2019) |
| p35S::Lti6b-GFP | Col-0 | (Kadota *et al.*, 2014) |

**Table S3:**
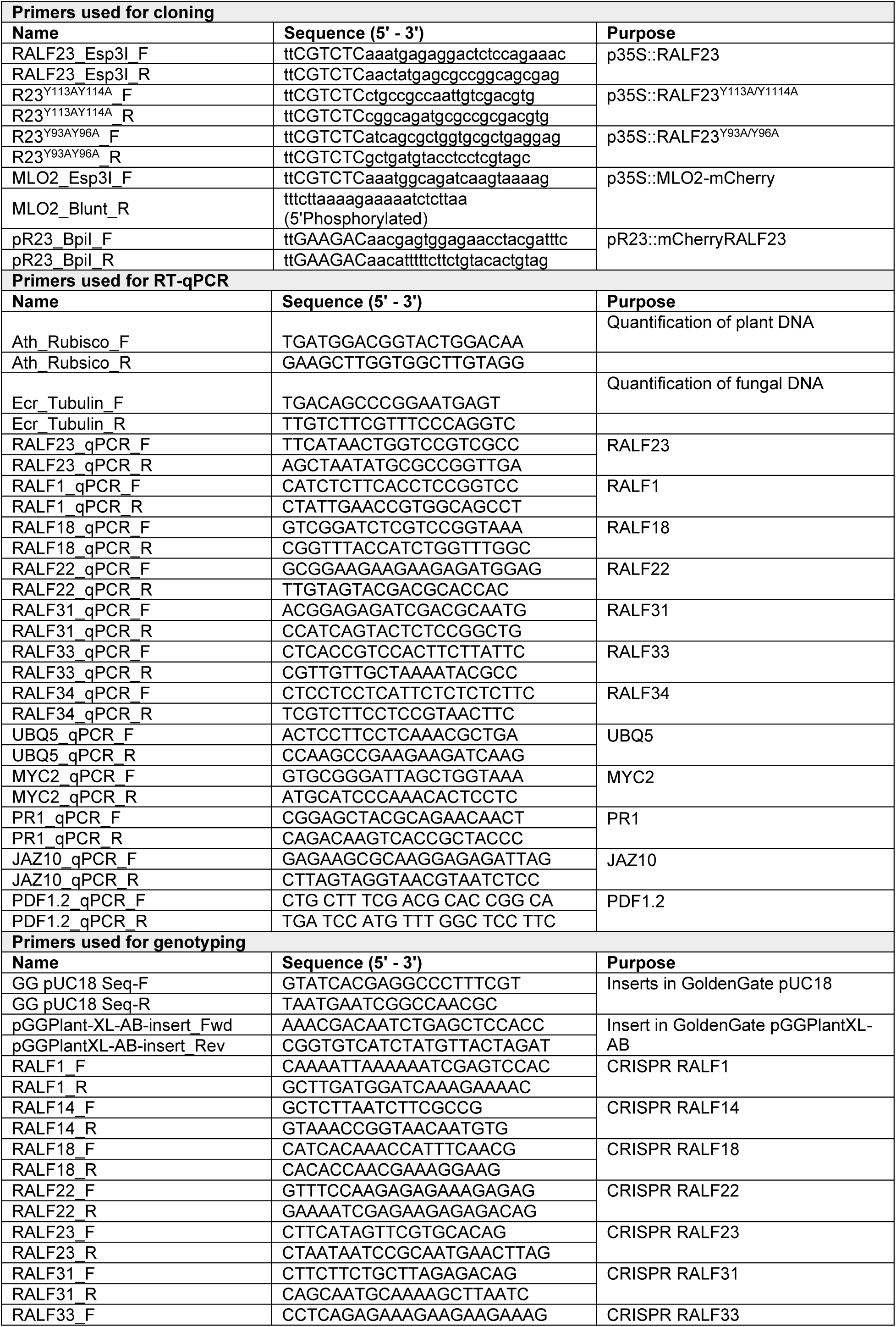
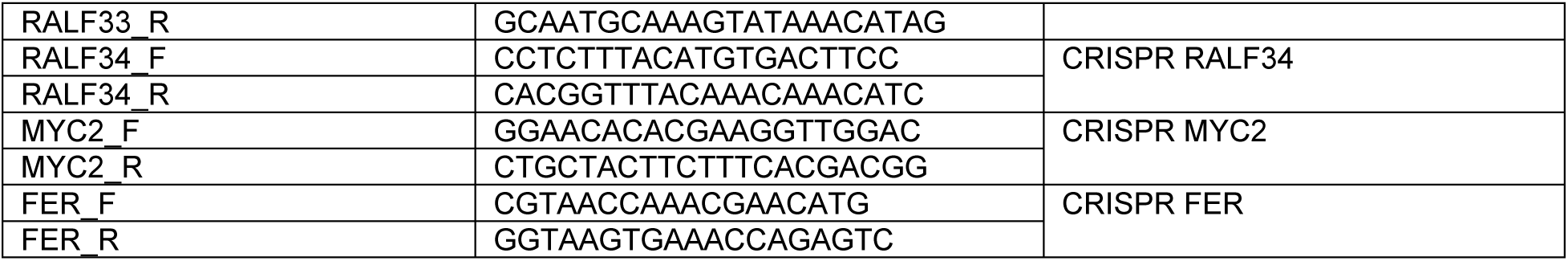
Primers used in this study.

**Table S4:**
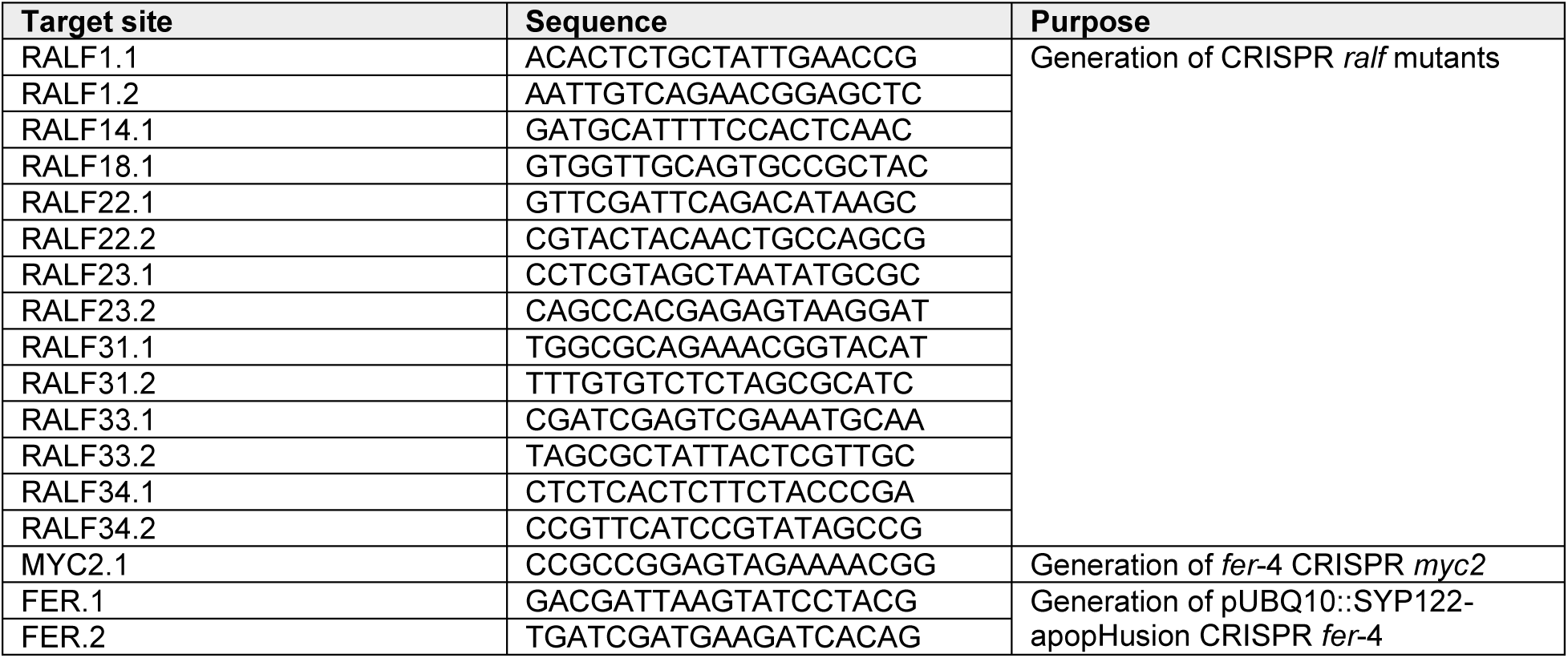
CRISPR target sites.

